# Screening of leaf extraction and storage conditions for eco-metabolomics studies

**DOI:** 10.1101/2023.06.26.546251

**Authors:** Jakob Lang, Sergio E. Ramos, Marharyta Smohunova, Laurent Bigler, Meredith C. Schuman

**Author notes:** **Author for correspondence:** Jakob Lang.

## Abstract

Mass spectrometry-based plant metabolomics is frequently used to identify novel natural products or study the effect of specific treatments on a plant’s metabolism. Reliable sample handling is required to avoid artifacts, which is why most protocols mandate shock freezing of plant tissue in liquid nitrogen and an uninterrupted cooling chain. However, the logistical challenges of this approach make it infeasible for many ecological studies. Especially for research in the tropics, permanent cooling poses a challenge, which is why many of those studies use dried leaf tissue instead. We screened a total of ten extraction and storage approaches for plant metabolites extracted from maize leaf tissue across two cropping seasons to develop a methodology for agroecological studies in logistically challenging tropical locations. All methods were evaluated based on changes in the metabolite profile across a 2-month storage period at different temperatures with the goal of reproducing the metabolite profile of the living plant as closely as possible. We show that our newly developed on-site liquid-liquid extraction protocol provides a good compromise between sample replicability, extraction efficiency, material logistics, and metabolite profile stability. We further discuss alternative methods which showed promising results and feasibility of on-site sample handling for field studies.

**Highlight:** We developed an on-site metabolite extraction method for leaf tissue samples from field studies in challenging logistical circumstances. We highlight extract stability and reproducibility compared to frozen or dried tissue.

## Introduction

In agriculture, high-throughput phenotyping approaches have become essential to assess traits related to increased yield, as well as those that confer tolerance to environmental stresses in crops (Araus and Cairns, 2014). Metabolomics is a powerful analytical approach that can provide information on the patterns and nature of plant responses to the environment, by providing information on the chemical features, identity, and quantity of metabolites produced by plants in different conditions (Sardans *et al*., 2021). In this way, metabolomics can add the chemical dimension to the high-throughput crop phenotyping toolbox, as thousands of metabolic markers often representing hundreds of metabolites can be recovered from a single leaf sample (Brunetti *et al*., 2013; Wolfender *et al*., 2015). Investigations of plant stress responses commonly focus on specialized metabolites, which are not essential for cell growth and development and are instead synthesized or modified by plants in response to specific environmental triggers (Macel *et al*., 2010; Walker *et al*., 2022).

Nevertheless, high-throughput phenotyping platforms have been developed under refined conditions (i.e., greenhouse and growth chamber facilities proximate to laboratories) and only reliably work with specialized equipment, which limits their application when dealing with realistic (field) conditions (Araus and Cairns, 2014). Such limitations extend to the use of a metabolomics approach in agriculture, where sample preparation and storage is a crucial step towards obtaining high quality data. For instance, most protocols in plant metabolomics require liquid nitrogen to shock-freeze the tissue immediately upon collection and keep the material frozen during the sample handling procedure. While this approach offers the closest representation of the metabolites in the living plant, it requires uninterrupted cooling (usually at −80 °C) and rapid sample handling to avoid thawing and degradation (Ossipov *et al*., 2008; Sedio *et al*., 2018; Bakhtiari *et al*., 2021).

A common alternative, when cooling conditions are not met, is to dry the plant tissue after collection and store the dried material, which is an attempt to stop enzymatic activity by removal of all water from the tissue. This approach would ideally be done by lyophilisation where the samples are completely frozen during the drying procedure, which should stop the enzymatic activity during the entire procedure (Walker *et al*., 2011). However, lyophilisers are usually only found in well-equipped laboratories and rarely available at field sites, which leaves drying in ovens (Fernandez-Conradi *et al*., 2022) or ambient conditions (Dela Cruz *et al*., 2022) as the main feasible alternatives, with desiccant supported drying as an alternative primarily established in DNA sequencing (Chase and Hills, 1991). The drying process allows for highly reproducible samples; however, little data is available on how the drying process changes the obtained metabolite profile due to differential stability of different metabolites. As a result, there is a need for a sample preparation method that ensures sample stability until the samples can be processed in the laboratory. This is particularly relevant when the sampling fields are located far from the laboratory facilities, and field campaigns are not easy or possible to repeat.

Here, we address limitations for the use of metabolomics in realistic agroecological conditions by describing and comparing sample handling methods. These methods were conceived in the context of a larger project aiming at understanding the metabolomic profile of maize grown under different conditions in tropical Africa, where weather and logistics conditions can make a metabolomics approach challenging. We first evaluated the suitability of two leaf preservation and six extraction methods, based on changes in metabolite profile across a 75-day storage period, to determine the method that resulted in the best apparent sample stability as judged by similarity to the metabolite profile obtained by standard laboratory procedures: solid-phase extraction (Glauser *et al*., 2011; Marti *et al*., 2013), or liquid-liquid extraction (Fiehn *et al*., 2000; Salem *et al*., 2016) of flash-frozen and finely powdered leaf tissue within a day after harvest. We then conducted a follow-up study focussing on an on-site liquid-liquid extraction procedure in comparison to in-field air-drying followed by laboratory extraction, and the laboratory standard procedure. Our results demonstrate that an on-site liquid-liquid extraction procedure generates reproducible metabolomic profiles while being feasible for field studies in terms of effort and stability of extracts. The methodology presented in this paper has the potential to be a viable alternative to the more established methods for plant metabolomics research in field studies and contribute to a better understanding of plant metabolism under realistic conditions (Peters *et al*., 2018).

## Materials and Methods

### Chemicals and materials

Acetonitrile (MeCN), methanol (MeOH) and isopropanol were obtained from *Biosolve* (ULC grade, Valkenswaard, Netherlands) and formic acid from *VWR Chemicals* (LC–MS grade, Dietikon, Switzerland). Ultrapure water (< 2 ppb TOC) was produced using a Milli-Q Advantage A10 water purification system (*Merck*, Burlington, MA, USA). For mass calibration, a 10 mM sodium formate solution was used, and ion mobility calibration was performed using ESI-L low concentration tune mix bought from *Agilent* (Santa Clara, CA, USA). The 10 mM sodium formate solution contained 1 M NaOH (250 μL) and formic acid (50 μL) in 50% isopropanol (25 mL). Dichloromethane (DCM) was purchased from *Honeywell* (Charlotte, NC, USA), Tween-20 from *Fisher Scientific* (Hampton, NH, USA) and all other chemicals from *Sigma-Aldrich* (St. Louis, MO, USA).

### Sample handling for broad method screening

Although we aim to develop a method practical for field research in tropical maize agroecosystems (i.e., central Africa), we required an experimental setting which allowed for comparison to extracts generated with an unbroken cooling chain. For this reason, maize plant tissue was collected from field-grown maize at the Strickhof Competence Centre of Agricultural Sciences (Eschikon, Switzerland, 47.4524090, 8.6806795) and used in eight different sample extraction and storage approaches. An overview of the employed methods is shown in **Fig. 1A** and a detailed description of all procedures can be found in the SI sections 1 and 2.

**Fig. 1:**
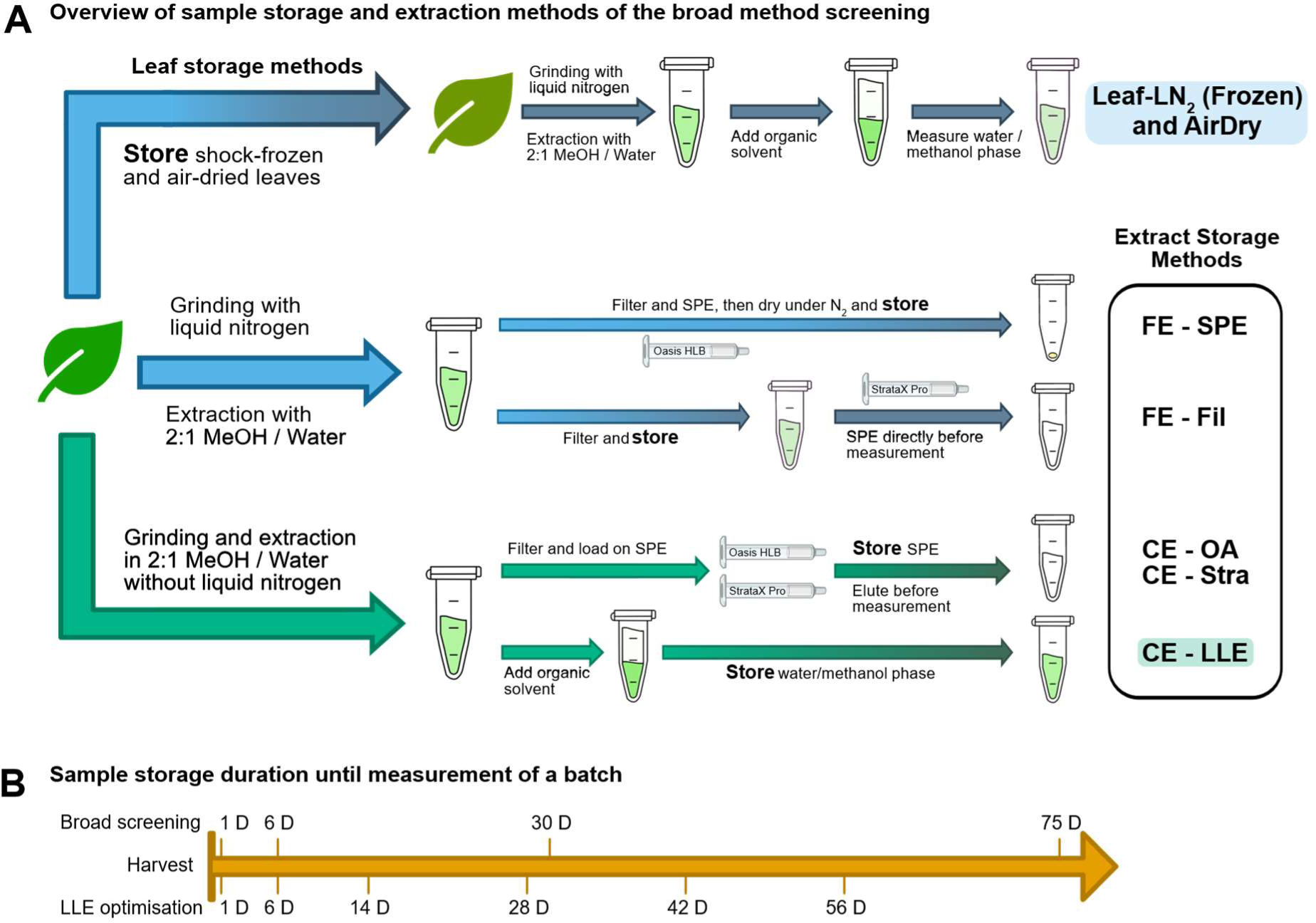
Overview of the evaluated sample extraction and storage methods (**A**). Blue arrows indicate extractions where liquid nitrogen was used during homogenisation (FE = Frozen Extraction), while green arrows indicate that no liquid nitrogen was used (CE = Crude Extraction). Bright colours indicate pre-storage processing, dark colours show sample preparation done after the storage period. Only the top pathway includes methods where leaf tissue is stored, either frozen or air-dried, the other pathways show the various leaf extract storage methods, which were prepared within 30 hours of harvest. The highlighted methods were later used during the LLE optimisation, where CE-LLE is referred to as “On-Site Extract storage”. The timeline (**B**) shows the evaluation time points of the broad method screening and the LLE Optimisation.

The samples were then stored at three different temperatures (30 °C, 4 °C, and −20 °C) for 1 day, 1 week, 1 month, and 75 days, respectively. At each of those timepoints four replicates of each method and of each storage temperature were analysed.

### Sample handling for liquid-liquid extraction optimisation

As a follow up study during the following cropping season, we evaluated metabolite stability in two extraction solutions and compared those results to air-dried and shock-frozen leaf storage. A detailed description of all procedures can be found in the SI sections 1 and 2. The samples were again stored at the same three different temperatures (30 °C, 4 °C, and −20 °C) and four replicates per timepoint, method and storage temperature were measured at six timepoints after 1 day to 8 weeks of storage time as shown in the timeline in **Fig. 1B**.

### UHPLC-HR-MS/MS Setup

Liquid chromatography was performed on a Vanquish Horizon UHPLC System by *Thermo Fisher* (Waltham, MA, USA) build from a Vanquish binary pump H, a Vanquish split sampler HT and a temperature-controllable Vanquish column compartment. Chromatographic separation was achieved on an ACQUITY Premier CSH C18 Column (130 Å, 1.7 µm, 2.1 × 50 mm, *Waters*, Milford, MA, USA) at 30 °C to reduce column backpressure. Eluent A consisted of H2O + 0.1% HCOOH and B of MeCN + 0.1% HCOOH. The solvent flow was kept at 0.6 mL/min with the following gradient: (i) 5% B isocratic from 0.0 to 0.4 min; (ii) linear increase to 35% B until 2.8 min; (iii) linear increase to 75% until 3.2 min; (iv) linear increase to 100% B until 3.3 min, (v) holding 100% B until 4.4 min (vi) back to the starting conditions of 5% B until 4.5 min; (vii) equilibration for 1.1 min until the next run. The injection volume is dependent on the employed extraction method and is specified in the detailed extraction protocols in SI sections 1 and 2.

A timsTOF Pro hybrid quadrupole-time-of-flight (QTOF) mass spectrometer equipped with trapped ion mobility spectrometry (TIMS) produced by *Bruker* (Bremen, Germany) was connected to the Vanquish UHPLC system and was used to acquire ion mobility and MS/MS data. Ionisation was performed in positive and negative ESI mode and the scan range was set to 20 to 1350 *m/z* at a 12 Hz acquisition rate. Mass and CSS calibration was performed using the *Agilent* low concentration tune mix (13 compounds in acetonitrile, part number G1969-85020) prior to analysis. For additional mass accuracy, a calibration segment was programmed from 0.05 to 0.15 min at every UHPLC run with the help of a 6-port-valve with a 20 µL loop which contained a solution of 10 mM sodium formate clusters.

### Software and Data Treatment

Instrument control was done using Hystar (*Bruker*, version 6.0) containing a Chromeleon Plug-In (*Thermo Fisher*, plugin version 1.3.8, Chromeleon version 7.3.0) and otofControl (*Bruker*, version 6.2). Data quality assessment was performed in DataAnalysis (*Bruker*, version 5.3) and data treatment (detailed below) in MetaboScape (*Bruker*, version 2022b). Figure plotting was done using python (version 3.8.5) in the Spyder IDE (version 5.0.3) using the libraries pandas (version 1.2.4), and bokeh (version 2.3.2). Posthoc analyses were performed with R (version 4.2.2) (Ihaka and Gentleman, 1996) with the library emmeans (version 1.8.3).

MetaboScape was used for peak picking, blank subtraction, data normalisation by internal standard, pareto transformation, and data evaluation with principal component analysis (PCA). The effects of pareto transformation were checked on representative datasets to ensure that this normalization and transformation resulted in a similar magnitude and approximately normal distribution of metabolite features across samples (Metaboanalyst (Pang *et al*., 2021), **Fig. S1 and S2**). All parameters for the peak picking and data evaluation are shown in the SI, section 3. The peak tables were exported in .csv format (see Data Availability) and PCA data was exported in .csv format to plot graphs using our python workflow (see SI, section 4). Compounds were classified with ClassyFire (Djoumbou Feunang *et al*., 2016), using InChi codes exported from MetaboScape.

### Recommended sample extraction procedure

For the full methods detailing all tested extraction procedures, see the detailed extraction protocols in SI sections 1 and 2. Here, we detail the recommended extraction procedure.

An extraction solution consisting of MeOH / water in a 2:1 ratio and camphorsulphonic acid as an internal standard (20 ng / mL) was prepared, of which 200 µL were added to a 1.5 mL Eppendorf tube for each sample. This solution is appropriate for extracting mid to high polarity metabolites which are commonly studied and contain many specialised secondary metabolites. Twelve leaf disks were collected with a 6 mm diameter hole punch (*Milian*, Vernier, Switzerland) directly into the extraction solution and the immersion in MeOH directly upon collection may reduce enzymatic activity in the sample (Maier *et al*., 2010). The tubes were thoroughly shaken and transported in a common household cooling box containing ice packs.

The leaf tissue was ground inside the Eppendorf tubes using plastic micropestles having a tip with approximately the same volume as the tip of the 1.5 mL Eppendorf tubes and attached to a household electric drill as shown in **Fig. S4**. It is recommended to use micropestles with a rough surface to facilitate leaf grinding, which we did by roughening the surface using 240 grit sandpaper. After the leaf tissue was ground to a paste, another 500 µL of the extraction solution was added before shaking thoroughly. The liquid-liquid extraction was performed through addition of 500 µL of chloroform to separate pigments and lipids, followed by thoroughly shaking. After letting the tubes rest for approximately 10 minutes at room temperature, the phase separation was completed, and 300 to 400 µL of the upper MeOH / water phase was transferred to fresh microcentrifuge tubes. For this study, grinding and liquid-liquid extraction was performed after transport of samples to a lab, but the procedure does not require any laboratory infrastructure and we have since performed it outside of laboratories for field studies (Lang *et al*., in preparation)

## Results

### Suitability of internal standards

For the broad method screening, we selected stevioside as an internal standard, but during the data evaluation we noted an issue, which led us to seek alternatives. In the mass spectrum of stevioside in **Fig. 2A**, the detected signals for the proton and ammonium adducts (805 and 822 *m/z*) are highlighted alongside the main signal at 319 *m/z* which matches the loss of all three hexose substructures. Additionally, signals were marked which match the loss of one and two hexose substructures.

**Fig. 2:**
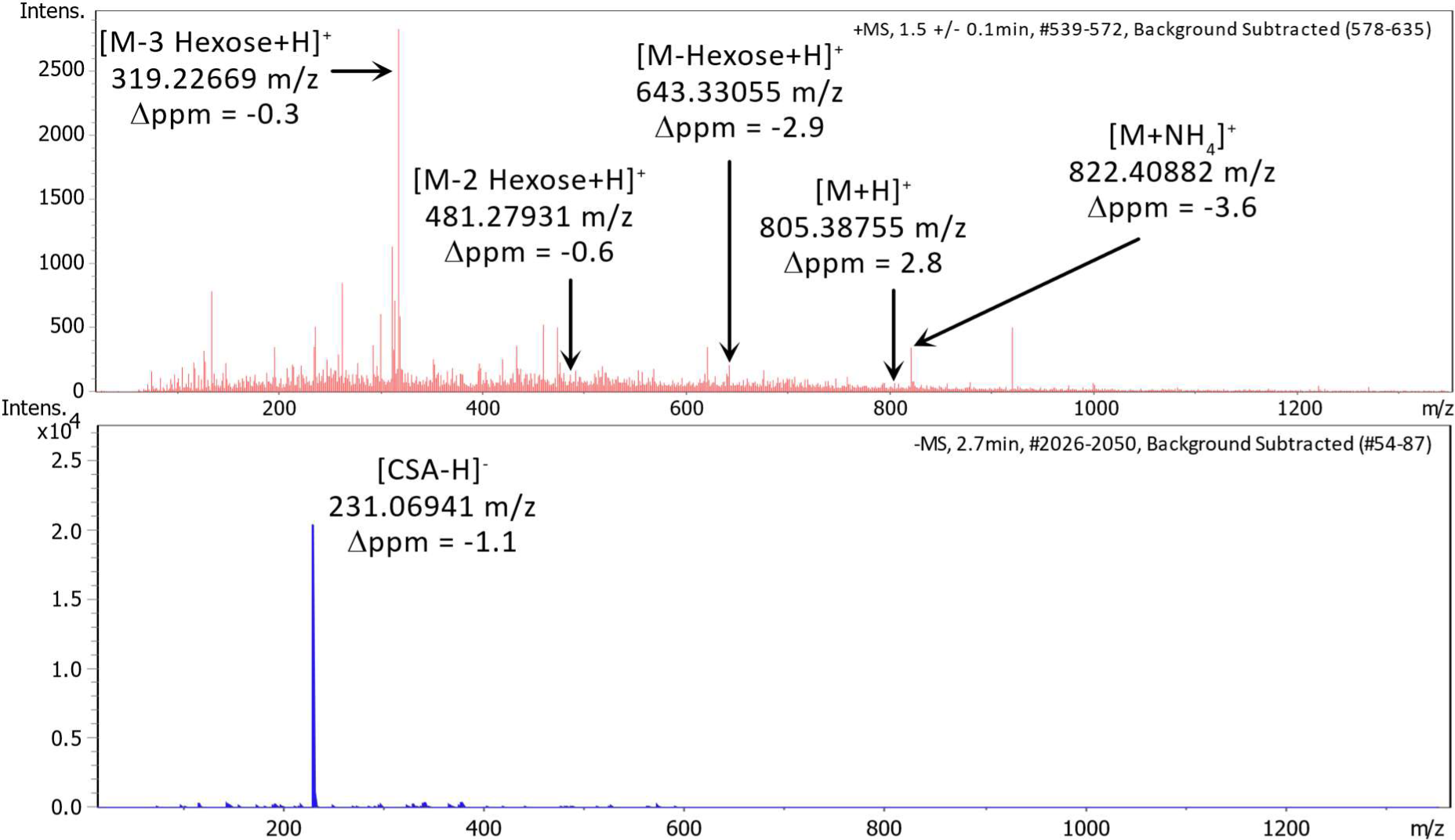
Comparison of the full scan MS spectrum of the internal standards stevioside (red, A) and camphorsulphonic acid (CSA, blue, B) with signal annotation of matching m/z ratios.

We attributed this to a possible in-source fragmentation and combined with a slight reduction in peak area observed with longer storage periods, the decision was made to include two additional possible internal standards – camphorsulphonic (CSA) and glycyrrhizic acid – in the LLE optimisation experiment. For comparison, the mass spectrum of CSA can be found below the stevioside spectrum in **Fig. 2B** and shows a single signal without any fragmentation. **Fig. S3** shows the intensity of each of the three compounds across the storage experiment. CSA showed a stable signal across the storage period with high ionisation efficiency, so we recommend using CSA over stevioside or glycyrrhizic acid. For targeted metabolomic analyses, isotopically labelled reference compounds would be preferable.

### Comparison of leaf homogenisation efficiency

Both during the broad method screening and later optimisation experiments, different approaches were tested for leaf tissue homogenisation using steel ball mills, ceramic mortars and micropestles. When freezing tissue in liquid nitrogen while grinding, a powder is generally obtained. However, when homogenizing air-dried leaf tissue with either ball mills or ceramic mortars, we were unable to obtain a powder, as some leaf veins remained intact. A direct comparison of the powders obtained when grinding fresh leaf tissue and air-dried leaf tissue in liquid nitrogen is shown in **Fig. 3**.

**Fig. 3:**
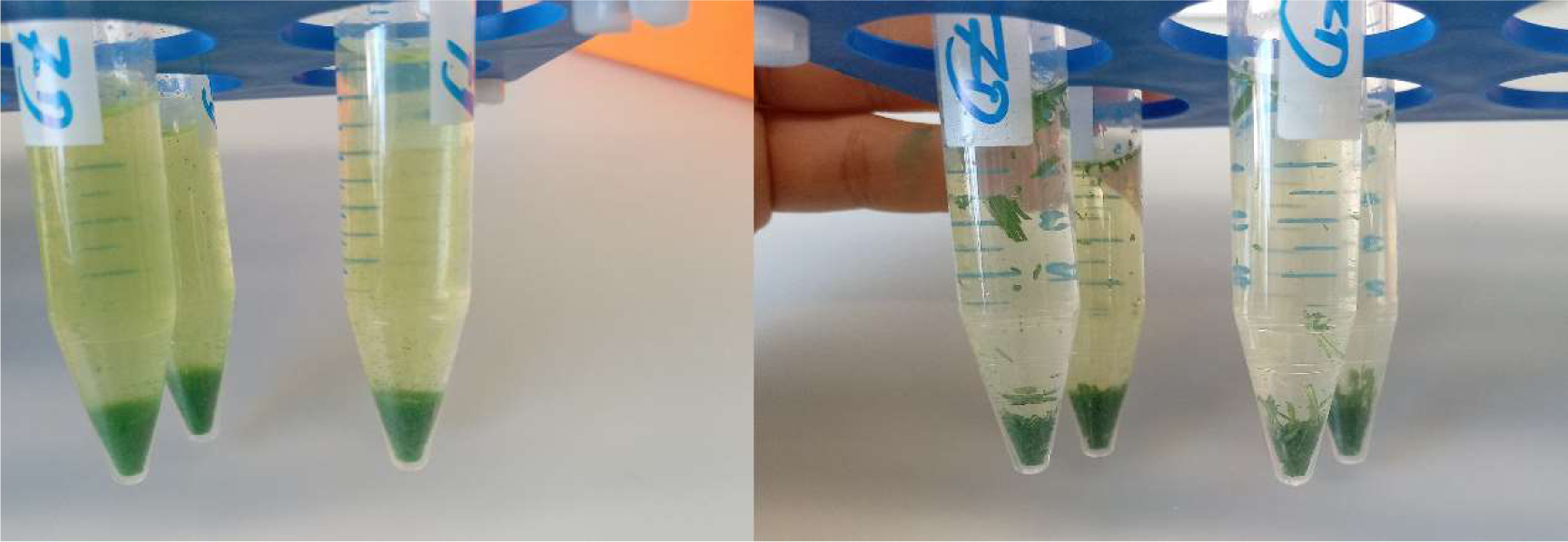
Comparison of ground flash-frozen (left) versus air-dried (right) leaf tissue following the same pulverization procedures.

Both of those methods still led to a more homogeneous product than attempting to grind tissue without using liquid nitrogen. Doing so with a ceramic mortar left the leaf tissue structure mostly intact, whereas with a micropestle, a chunky and more homogeneous paste could be obtained (**Fig. S4**).

### Selectivity of sample preparation methods

During the broad method screening, fundamentally different sample purification approaches were tested, most notably solid-phase extraction (SPE) and liquid-liquid extraction (LLE). The two approaches lead to significant differences in the resulting metabolite profile. In our experiments, the profile after sample workup with SPE was shifted towards molecules with a higher molar mass and a lower polarity compared to samples prepared by LLE, which is to be expected based on the fundamental selectivity of the methods. The highest polarity compounds are lost while washing the SPE cartridge with water, while lipids and other low polarity compounds are later eluted with MeOH together with the polar metabolites. Comparing this to LLE, higher polarity compounds including salts are retained in the water/methanol phase, while lower polarity compounds are lost in the organic phase. This trend can already be observed in a base peak chromatogram, as shown in **Fig. 4A and can** further be explored when comparing the compound classes that could be identified. The key difference between the methods is the large gap in the number of identified organic acids which are mostly absent in samples extracted by SPE as highlighted in **Fig. 4B**. Notably we did not perform an annotation with a lipid specific spectral database, which likely would highlight a larger annotation rate in the SPE samples.

**Fig. 4:**
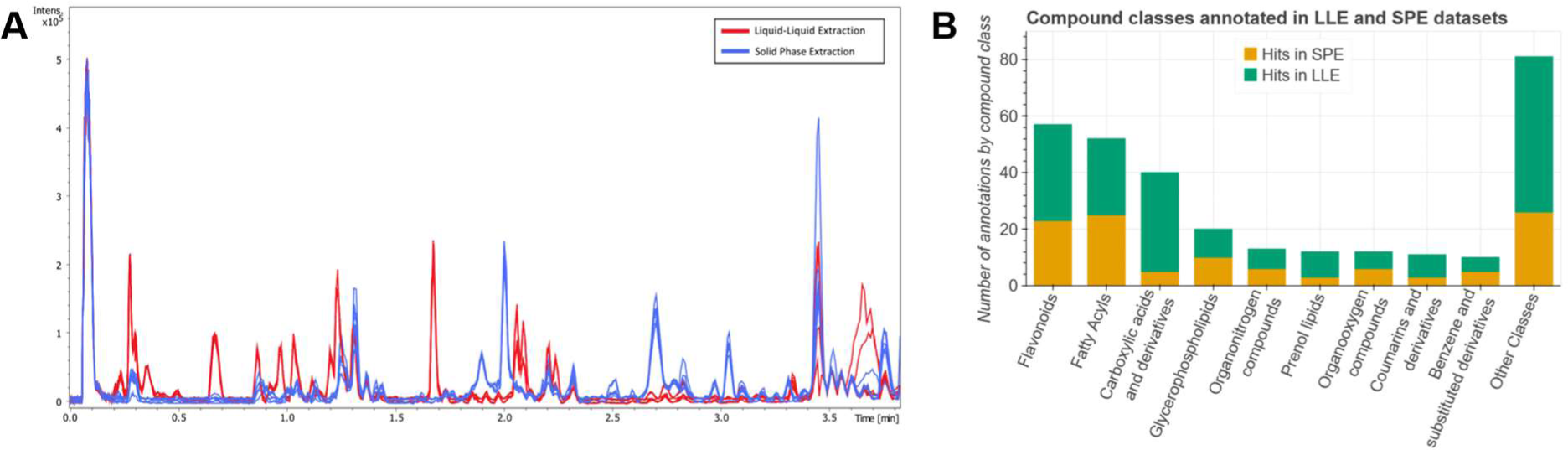
Overlaid chromatograms of a subset of four samples prepared by solid-phase extraction (SPE, blue) and four samples prepared by liquid-liquid extraction (LLE, red) highlighting generally higher abundance of high-polarity (shorter retention time) compounds in LLE samples (**A**) and a comparison of annotated features by compound class which further highlights the different extraction efficiencies (**B**).

The significant shift of the metabolite profile causes a challenge when it comes to multivariate data comparison, where principal component analysis (PCA) is a common approach. Any PCA which contains LLE and SPE samples will group the extraction approaches tightly together as shown in **Fig. S5**, which masks the shifts in the profile across a storage period. Thus, all PCA results were plotted separately for LLE and SPE sample groups (**Fig. S6** and **S7**) to allow a sensible interpretation.

### Extract stability over time

Changes in the overall metabolite profile were assessed by PCA, which showed that in almost all cases the metabolite profile changed the most when samples were stored at 30 °C (listed as room temperature, RT). During the broad method screening, the metabolite profile continued to shift for all evaluated sampling methods (**Fig. S8** to **S13**) without reaching a stable result (which could occur after completing all possible molecular transformations). Examples of the PCA can be found in the SI section 6 with special attention towards **Fig. S6** and **S7**, which show the comparison of all evaluated LLE and SPE methods. During the LLE optimisation experiment, the shift of the metabolite profile over time was significantly reduced. As an example, **Fig. 5C** shows the PCA of all samples prepared using the on-site sample extraction procedure across, including all storage temperatures and timepoints. Notably, samples stored at room temperature are shifted along PC1 with longer storage duration shifting to higher PC1 values, while cooled samples (both 4 °C and −20 °C) cluster tightly together with smaller PC1. A minor trend towards higher PC1 values can be seen for samples stored at 4°C. When excluding the room temperature samples, all datapoints cluster randomly in PC1 and 2 (**Fig. S17**), while higher PC dimensions show a minor shift over time, which is highlighted in **Fig. 5E**. For comparison, results from storing shock-frozen leaf tissue at −20 °C are shown in **Fig. 5D** and demonstrate a shift of the metabolite profile along PC2.

**Fig. 5:**
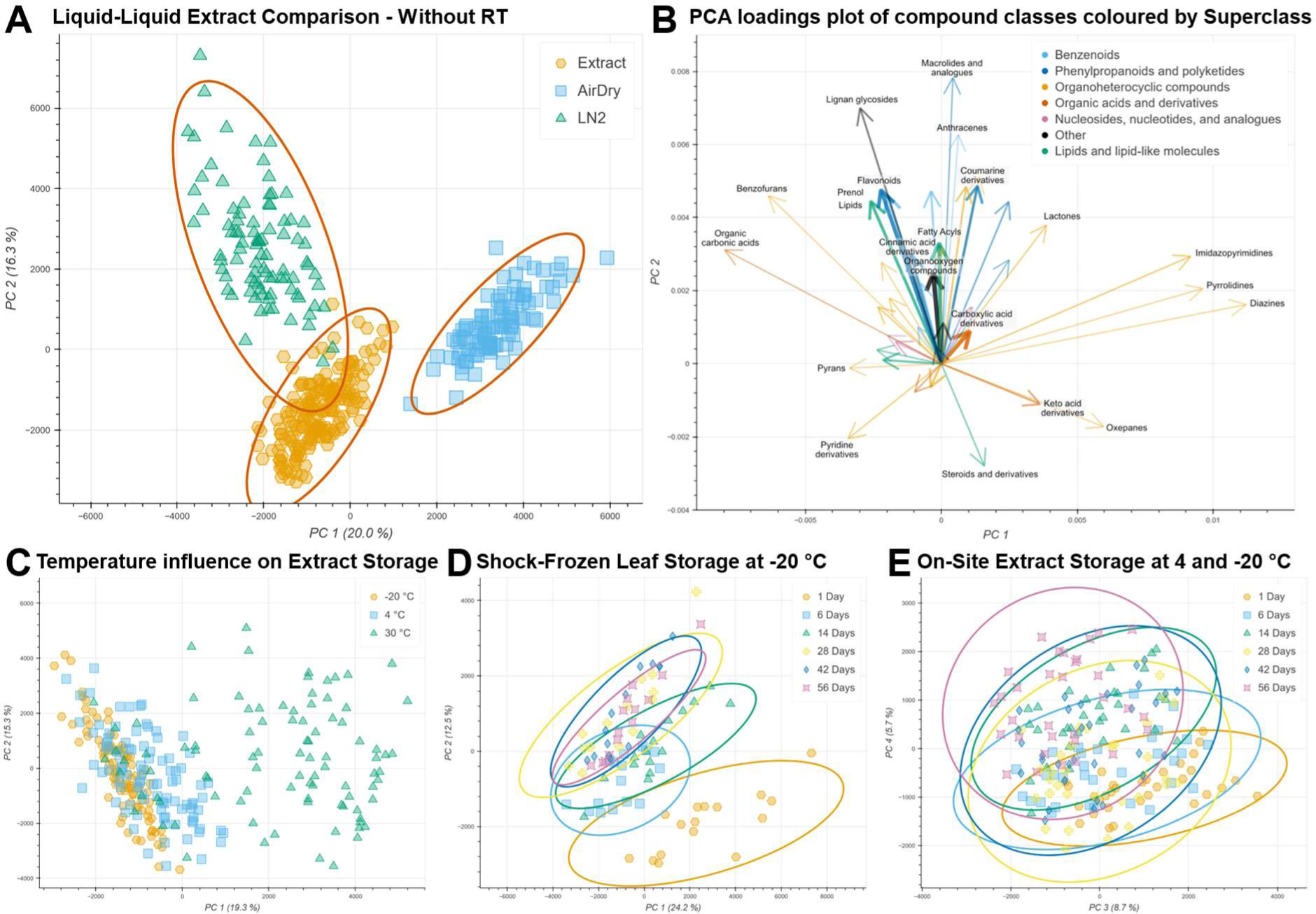
Principal components analysis (PCA) conducted on metabolite profiles of samples extracted and stored under different conditions: (A) LLE of differently handled leaf tissue samples, (B) compound class impact on the separation of 5A, arrow width indicates number of compounds of each class in a range between 5 and 199 compounds, (C) influence of storage temperature on the metabolite profile shift of LLE extracts (includes all storage timepoints), (D) profile shift over time for frozen storage of shock-frozen leaf tissue, (E) shift of on-site LLE of both −20 °C and 4 °C.

The shifts of the metabolite profile over the storage period can also be seen in the direct comparison of the three storage methods in **Fig. 5A**, where larger shifts of the metabolite profile led to a wider distribution across the PC dimensions. The shock-frozen leaf tissue shows the widest spread of all methods, primarily in the direction of PC2 (see also **Fig. S15**), while both the air-dried leaf tissue (see also **Fig. S18**) and the on-site extraction samples show a much tighter grouping, indicating a more stable metabolite profile over the storage duration. The samples from air-dried leaf tissue are fully separated from the other methods along PC1 while the on-site extraction is separated from shock-frozen leaf tissue samples along PC2 with an overlapping 95% confidence ellipse. Those trends are shown under the exclusion of samples stored at room temperature for clearer grouping of replicates but can also be observed when including those samples as shown in **Fig. S14**.

The separation of sample storage methods is influenced by various compound classes as seen in the merged loadings plot in **Fig. 5B & Fig. S19**. Of the most abundant compound classes, the clearest trend emerges for flavonoids, which indicates an increased abundance in shock-frozen leaf samples with a short storage duration. Other frequently detected compound classes such as fatty acyls, cinnamic acid derivatives, and prenol lipids show similar trends and of all classes with 50+ annotated signals, only carboxylic acids show a minor trend to positive PC2 values, which is where air-dried samples are grouped. The strong shift of the metabolite profile of air-dried leaf storage samples can also be seen when comparing the identified compound classes of the three methods as shown in **Fig. 6**. Multiple compound classes, such as carboxylic acids and coumarin derivatives, show a reduced annotation count in the air-dried dataset, while shock-frozen and on-site extracts show comparable annotation rates for most compound classes.

**Fig. 6:**
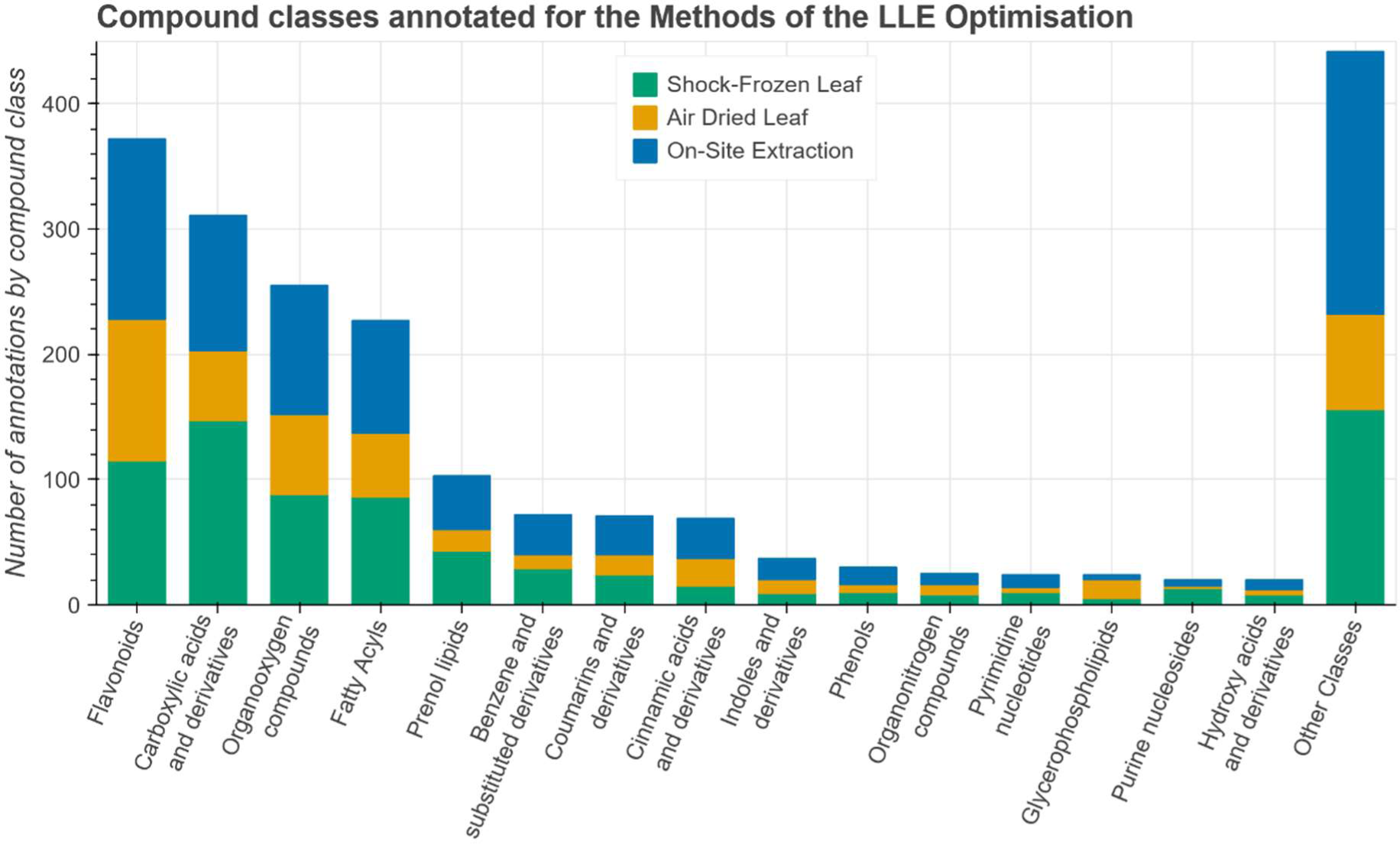
Annotated compound classes of the three methods tested during the LLE optimisation.

Lastly, a MANOVA analysis was performed which showed significant differences based on the storage method in PC 1 to PC 5 and a follow-up Tukey pairwise comparison based on the first five PCs (see: SI, section 5, posthoc analysis and **Fig. S20** and **S21**) indicated that all three groups are significantly different from each other. The comparison of the on-site extract storage and shock-frozen leaf storage samples showed the lowest degree of significance with a p-value of 0.0006, while p-values of any comparison involving the air-dried leaf storage samples were too small to be fully calculated (below 0.0001).

We attempted to show the effect of storage on plant stress biomarkers by inducing the maize plants with methyl jasmonate a day before sample collection. However, for the LLE optimisation experiment the plants were sown out earlier, which meant plants were already 14 weeks old at the time of sampling. That late in their development the reaction to stressors is reduced (Çakir, 2004) and we were thus unable to determine clear differences between stressed and unstressed plants as seen in **Fig. S16**.

### A note on storage of extracts on SPE cartridges

During the broad method screening we found indications that metabolite storage on solid-phase extraction (SPE) cartridges (procedure CE-OA in SI, section 1) could be a viable alternative for on-site sample preparation and storage. **Fig. 7** shows the samples stored on the SPE cartridge in comparison to samples that were dried under nitrogen flow after the SPE, storing the dried residue (procedure FE-SPE in SI, section 1). The samples stored on the cartridge seemed more reproducible (tighter grouping of replicates) and with a comparable shift over time compare to the samples following the “FE-SPE” procedure. If the focus of a study is on lower polarity and higher mass compounds, this method might be preferable to an LLE-based approach. However, due to material shortages at the time, the “CE-OA” approach was only evaluated at three storage timepoints, and we would therefore recommend more in-depth testing before employing this approach on a larger scale.

**Fig. 7:**
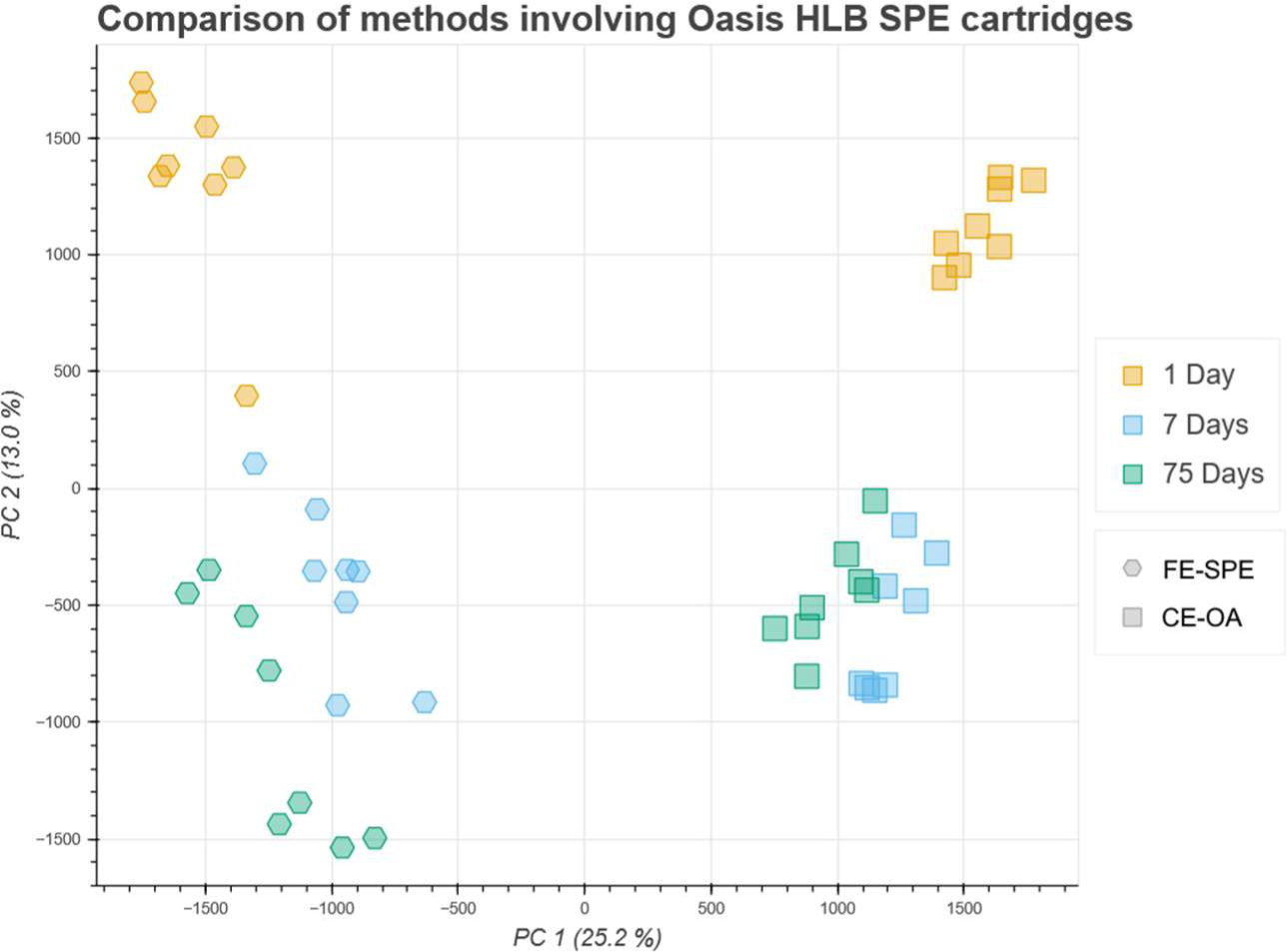
Principal component analysis of samples stored on an SPE cartridge (CE-OA, squares) and samples prepared by SPE and dried down for storage (FE-SPE, hexagons). Samples stored at 30 °C are not included and FE-SPE samples stored for 28 days were removed as there was no CE-OA counterpart for the direct comparison.

## Discussion

### Liquid-liquid extraction – extract and leaf storage

Storage of samples after an LLE without shock-freezing of the leaf tissue showed promising results during the broad method screening. All samples from the 7- and 30-day timepoints that were stored at reduced temperatures were tightly grouped together on the PCA and the samples stored at 4 °C and −20 °C showed a comparable metabolite profile (**Fig. S10**), which led us to study the LLE approaches in more detail. During the LLE optimisation experiment we could verify the minimal impact of storage in a freezer compared to refrigerator and obtained a highly reproducible metabolite profiles for both conditions (**Fig. 5A** and **E**). Overall, our on-site extraction procedure results in samples which more closely represent the metabolite profile of shock-frozen leaf tissue compared to air-dried leaf storage as seen in **Fig. 5A**. Additionally, the compound class analysis shown in **Fig. 6** was able to provide similar annotation rates for shock-frozen leaf tissue storage and the on-site extraction procedure. Even when including the samples stored at room temperature, the profile is closer to our goal than air-dried samples (**Fig. S14**), but there is a notable change depending on storage duration. As such, the storage duration of each sample would become an important factor to control for, which may not be required when storing the extracts at reduced temperatures.

**Fig. 5D** and **E** highlight the extract stability over storage duration, and notably a lower overall change in the metabolite profile than storage of shock-frozen leaf tissue. The metabolite profile of samples from air-dried leaf tissue is also very stable over the storage duration once the drying process is completed (**Fig. 5A** and **S18**), but multiple compound classes are no longer detected in dried leaf tissue as seen in **Fig. 6**. The minor shifts of the metabolite profile of both air-dried leaf and on-site extract storage allows the comparison of samples even if the storage duration is not the same across the dataset, which is not a given for shock-frozen leaves stored at −20 °C. The low rate of change over the storage duration of the on-site extracts might be related to the fact that all leaf material is collected into tubes that already contain 200 µL of the extraction solution, which consists of two thirds MeOH and one third water. MeOH has been shown to quench enzymatic activity and is frequently used before metabolite extraction from microbial extracts (Faijes *et al*., 2007; Link *et al*., 2008). We thus hypothesize that the immediate contact with MeOH assists with quenching of enzymatic activity for leaf tissue, not unlike flash-freezing with liquid nitrogen. The stability of the MeOH-immersed leaf tissue then becomes relatively independent of temperature and handling. Drying leaf tissue for storage and transport does not have such a quenching step after collection and drying takes more time than flash-freezing or penetration of leaf discs by MeOH solution. Similar effects have been described previously (Maier *et al*., 2010) and the instantaneous contact to the solvent seems to be a common theme to assure sample reproducibility.

### SPE as a potential candidate for lower polarity metabolites

We found that storage of extracts on SPE cartridges seemed to result in reproducible metabolite profiles across storage times and conditions, albeit with lower replication than for the other methods tested in broad method screening, due to material shortages at the time the work was conducted. As outlined before, SPE shows a significant difference in the metabolite profile compared to LLE and thus may be better suited for research focussing on lower polarity compounds (Šimura *et al*., 2018). As our aim was to find a method that can be applied for field studies, the additional logistical challenge of operating a vacuum pump to load the extract onto an SPE cartridge was deemed too large of a hurdle and we proceeded with a focus on LLE-based approaches instead. Besides the operation of a vacuum system, an additional downside is the increased material cost and logistics, which we estimate to at least double the cost per sample.

### Feasibility for field studies

While there are well established procedures for metabolomics sample handling under controlled conditions – most relying on shock freezing in liquid nitrogen followed by uninterrupted cooling to −80 °C (Balmer *et al*., 2013) – this approach is challenging to apply in field studies. A commonly used approach is to dry the plant tissue (ElNaker *et al*., 2021), which allows for reproducible results without any cooling; however, the metabolite profile is significantly impacted by the drying process, as shown by the significant separation along PC1 in **Fig. 5A**. Our proposed on-site LLE protocol, where a liquid extract is stored in commercial refrigerators, can help fill the gap between shock-frozen and dried leaf extracts. The sample extraction requires some low-cost laboratory chemicals and consumables and almost no infrastructure. Access to electricity is required for the drill for leaf homogenisation (at least to charge a battery) and a refrigerator allows for sample storage over at least two months with minimal shifts in the metabolite profile. While these requirements entail greater logistical challenges than the commonly used dried plant material method, avoiding the drying process can be worthwhile, especially if more labile metabolites are a focus of the study (Wu *et al*., 2023).

### Limitations of the proposed approach

While the on-site liquid-liquid sample extraction is feasible under logistically challenging conditions and provides samples which more closely represent the metabolite profile obtained from shock-frozen leaves than air-dried leaf storage, it comes with various limitations to consider before using it in large-scale field studies. Extracting metabolites on-site is a time-consuming task which requires some practice before employment in the field. Especially the tissue homogenisation can lead to significant variation between samples until a certain level of practice is reached. Since the exact degree of homogenisation is challenging to standardise, it is also ideally done by one person only to avoid person-to-person variations (Creydt *et al*., 2018).

The main limitation is that none of the evaluated methods was able to fully reproduce the metabolite profile obtained from shock-frozen leaf tissue with immediate sample processing. Any storage period did introduce significant shifts in the metabolite profile, even storing shock-frozen leaves at −20 °C. Whether the shifts of the metabolite profile are relevant for a specific application depends on the exact compounds of interest and cannot be generalised here. Furthermore, metabolite analyses often attempt an uninterrupted cooling chain at −80 °C, which is common in greenhouse experiments, but is a significant logistical challenge for field studies (Nagler *et al*., 2018). Our dataset did not include leaf storage at −80 °C which might lead to a reduced shift of the metabolite profile compared to storage at −20 °C. Lastly, the on-site LLE method for sample collection and extraction was thoroughly tested on maize plants, but no other species was used during this study. Since specialised metabolites of other plants can show a different degradation behaviour, the procedure might not be suitable for all plant metabolomics studies.

## Supplementary Material

Section 1: Detailed description of the broad method screening methods

Section 2: Detailed description of the LLE optimisation methods

Section 3: Detailed description of the data evaluation procedure

Section 4: Python script for plotting of PCA data

Section 5: R script and output of MANOVA and Pairwise comparison

Section 6: Additional pictures and graphs

- Fig. S1 and S2: Effects of pareto scaling on the LLE optimisation dataset.
- Fig. S3: Signal of the three internal standard candidates depending on storage time.
- Fig. S4: Photos of the on-site leaf homogenisation and extraction procedure
- Fig. S5 – S13: Additional PCA plots of the broad method screening.
- Fig. S14 – S21: Additional PCA plots of the LLE optimisation.

## Abbreviations

CE: Crude Extract (homogenisation at room temperature)
CSA: Camphorsulphonic acid
DCM: Dichloromethane
FE: Frozen Extract (homogenisation using liquid nitrogen to freeze the leaf)
LLE: Liquid-liquid extraction
MeCN: Acetonitrile
MeOH: Methanol
OA: Oasis HLB branded SPE cartridges
PCA: Principal component analysis
SPE: Solid-phase extraction

## Acknowledgments

We thank Marco Landis and other staff members of the Strickhof (Lindau, Eschikon, Switzerland) for allowing us to collect samples in their maize fields. We thank Urs Stalder and Karoline Rehm for support with maintenance and method development on the UHPLC-MS system.

## Author Contributions

JL, MCS: conceptualization; JL: data curation; JL, SER: formal analysis; LB, MCS: funding acquisition; JL, SER, MS, MCS: investigation; JL, MCS: methodology; JL, MCS: project administration; LB, MCS: Resources; JL, SER: Software; LB, MCS: Supervision; JL, MCS: Validation; JL: Visualization; JL, SER, MCS: Writing – original draft; Writing – review & editing: JL, SER, MS, LB, MCS.

## Conflict of interest

No conflict of interest declared.

## Funding

This work was supported by the European Research Council (ERC) under the European Union’s Horizon 2020 research and innovation programme under the grant agreement UPSCALE (SFS-35-2019-2020).

## Data availability

Processed peak tables and MS/MS fragmentation patters are available on Zenodo (Lang *et al*., 2023): https://zenodo.org/doi/10.5281/zenodo.10219180.

## Notes

### Competing Interest Statement

The authors have declared no competing interest.

### Summary of Updates

Most parts of the publication (except the experimental section) were revised to clarify some questions. The datasets were reanalysed to add compound classifications to all annotated signals, which was used to create new figures (4B, 5B, and 6) to better illustrate key differences among the extraction and storage methods. Most figures were revised for clarity and the introduction and discussion were expanded with additional context.

