## Supplementary Information for "Screening of leaf extraction and storage conditions for eco-metabolomics studies"

- Supplementary methods 1 & 2: Extraction and Measurement Methods
- Supplementary methods 3: MetaboScape Data Handling
- Supplementary methods 4 & 5: Python and R Scripts for plotting and MANOVA
- Supplementary figures 1-21
- References

#### 1. Broad Screening Methods

##### 1.1. Leaf Tissue Collection 2021

Maize plants of variety LG 31272 were grown under field conditions at the Strickhof competence centre in agricultural sciences (Eschikon, Switzerland, 47.4524090, 8.6806795) and tissue collection took place seven weeks after seeding on 29th July 2021. From all sampled maize leaves, the tip (roughly 10 cm) was cut off before collecting samples as leaf strips of 2 cm width with approximately 10 strips being collected per leaf. The leaf strips were split into three groups and subjected to different sample handling and extraction procedures. One group was packed in envelopes and left to dry at ambient temperature (approximately 26 °C), while the other two groups were packed in aluminium foil and either stored on ice, or shock-frozen in liquid nitrogen before being stored on dry ice.

#### 1.2. Sample Preparation

##### 1.2.1. Unprocessed Leaf Storage (Leaf-LN2)

The leaf strip storage used strips of all three transport handling groups. Leaves stored at -20 °C came from the group that was shock frozen and stored on dry ice, leaves stored at 4 °C were kept on ice during transport, and leaves stored at 30 °C were transported at ambient temperature.

Before the measurement, leaf squares of 2-by-2 cm were ground with a ceramic mortar and pestle while covered with liquid nitrogen. The ground leaf powder was then extracted using 5 mL per sample of a MeOH/$H_2O$ mixture in 2:1 ratio, containing 130 ng/µL of stevioside as internal standard (but see main text for discussion of preferred internal standards) and acidified with 0.1 % formic acid. The solution was shaken thoroughly and rested at room temperature for 30 minutes before filtering through a filter paper (*Macherey-Nagel*, Type 615). A liquid-liquid extraction was performed using 1 mL of the extract and 500 µL of dichloromethane (DCM). The upper phase was transferred to a vial and used for measurement with a 3 µL injection volume.

##### 1.2.2. Air-dried Leaf storage (AirDry)

This extraction used leaf strips that were dried at ambient temperature after collection. Since the leaves were not fully dried when reaching the laboratory, the strips were stored at 30 °C overnight to fully dry the tissue before storing them at -20 °C, 4 °C or 30 °C.

Before the measurement, leaf squares of 2-by-2 cm were ground with a ceramic mortar and pestle while covered with liquid nitrogen. The ground leaf powder was then extracted using 5 mL per sample of a MeOH/$H_2O$ mixture in 2:1 ratio, containing 130 ng/µL of stevioside as internal standard (but see main text for discussion of preferred internal standards) and acidified with 0.1 % formic acid. The solution was shaken thoroughly and rested at room temperature for 30 minutes before filtering through a filter paper (*Macherey-Nagel*, Type 615). A liquid-liquid extraction was performed using 1 mL of the extract and 500 µL of dichloromethane. The upper phase was transferred to a vial and used for measurement with a 3 µL injection volume.

###### 1.2.3. Freeze Extract – Post SPE Storage(FE-SPE)

This extraction used leaf strips that were shock-frozen and transported on dry ice. For each sample, a 2-by-2 cm square of leaf tissue was used. The squares were ground with a ceramic mortar and pestle while covered by liquid nitrogen. The ground leaf powder was then extracted using 5 mL per sample of a MeOH/$H_2O$ mixture in a 2:1 ratio, containing 130 ng/µL of stevioside as internal standard (but see main text for discussion of preferred internal standards) and acidified with 0.1 % formic acid. The solution was shaken thoroughly and rested at room temperature (approx. 25 °C) for 30 minutes before filtering through a filter paper (*Macherey-Nagel*, Type 615). The extract was purified by solid phase extraction on Oasis PriME HLB cartridges (*Waters*, Milford, MA, USA) containing 30 mg sorbent. The cartridge was loaded without prior conditioning with 500 µL of filtered plant extract and then washed with 1 mL of $H_2O$. The samples were then eluted with 2x500 µL of MeOH+0.1% formic acid. The solvent was then evaporated under nitrogen flow at 30 °C until dried completely and the dried samples were stored at -20 °C, 4 °C and 30 °C, respectively.

Before measurement, the samples were reconstituted in 1 mL of MeOH+0.1% formic acid and 3 µL were used as injection volume.

###### 1.2.4. Freeze Extract – Filtrate Storage (FE-Fil)

This extraction used leaf strips that were shock-frozen and transported on dry ice. For each sample, a 2-by-2 cm square of leaf tissue was used. The squares were ground with a ceramic mortar and pestle while covered by liquid nitrogen. The ground leaf powder was then extracted using 5 mL per sample of a MeOH/$H_2O$ mixture in a 2:1 ratio, containing 130 ng/µL of stevioside as internal standard (but see main text for discussion of preferred internal standards) and acidified with 0.1 % formic acid. The solution was shaken thoroughly and rested at room temperature (approx. 25 °C) for 30 minutes before filtering through a filter paper (*Macherey-Nagel*, Type 615). Replicate extracts were stored at -20 °C, 4 °C and 30 °C.

Before the measurement, the samples were purified by solid phase extraction on Strata X Pro cartridges (*Phenomenex*, Torrance, CA, USA) containing 30 mg sorbent. The cartridge was loaded without prior conditioning with 500 µL of filtered plant extract and then washed with

1 mL of $H_2O$. The samples were then eluted with 2x500 µL of MeOH+0.1% formic acid and the elute was used directly for measurement with a 3 µL injection volume.

###### 1.2.5. Crude Extract – Loaded Oasis SPE Storage (CE-OA)

This extraction used leaf strips that were transported on ice after collection. For each sample a 2-by-2 cm square of leaf tissue was used. The squares were ground with a ceramic mortar and pestle without addition of liquid nitrogen. The mashed leaf paste was then extracted using 5 mL per sample of a MeOH/$H_2O$ mixture in 2:1 ratio containing 130 ng/µL of stevioside as internal standard (but see main text for discussion of preferred internal standards). The solution was shaken thoroughly and rested at room temperature (approx. 25 °C) for 60 minutes before filtering through a filter paper (*Macherey-Nagel*, Type 615). An Oasis PRiME HLB cartridge (*Waters*, Milford, MA, USA) containing 30 mg sorbent was then loaded with 500 µL of the extract. After running the SPE cartridge dry, the replicate cartridges were stored at -20 °C, 4 °C or 30 °C.

Before the measurement, SPE cartridges were eluted with 2x500 µL of MeOH+0.1% formic acid and the elute was used directly for measurement with 3 µL injection volume.

###### 1.2.6. Crude Extract – Loaded StrataX SPE Storage (CE-Stra)

This extraction used leaf strips that were transported on ice after collection. For each sample a 2-by-2 cm square of leaf tissue was used. The squares were ground with a ceramic mortar and pestle without addition of liquid nitrogen. The mashed leaf paste was then extracted using 5 mL per sample of a MeOH/$H_2O$ mixture in 2:1 ratio containing 130 ng/µL of stevioside as internal standard (but see main text for discussion of preferred internal standards). The solution was shaken thoroughly and rested at room temperature (approx. 25 °C) for 60 minutes before filtering through a filter paper (*Macherey-Nagel*, Type 615). A Strata X Pro cartridge (*Phenomenex*, Torrance, CA, USA) containing 30 mg sorbent was then loaded with 500 µL of the extract. After running the SPE cartridge dry, the replicate cartridges were stored at -20 °C, 4 °C or 30 °C.

Before the measurement, SPE cartridges were eluted with 2x500 µL of MeOH+0.1% formic acid and the elute was used directly for measurement with 3 µL injection volume.

##### 1.2.7. Crude Extract – Liquid-Liquid Extract Storage (CE-LLE)

This extraction used leaf strips that were transported on ice after collection. For each sample a 2-by-2 cm square of leaf tissue was used. The squares were ground with a ceramic mortar and pestle without addition of liquid nitrogen. The mashed leaf paste was then extracted using 5 mL per sample of a MeOH/$H_2O$ mixture in 2:1 ratio containing 130 ng/µL of stevioside as internal standard (but see main text for discussion of preferred internal standards). The solution was shaken thoroughly and rested at room temperature (approx. 25 °C) for 60 minutes before filtering through a filter paper (*Macherey-Nagel*, Type 615). A liquid-liquid extraction was done by mixing DCM with the obtained leaf extract in 1:2 ratio. The solution was thoroughly shaken, and 1 mL of the upper phase was transferred into a vial before storage at -20 °C, 4 °C or 30 °C. The solution was used for measurement using 3 µL injection volume.

##### 1.2.8. Crude Extract – Silicone Tubing Storage (CE-ST)

This extraction used leaf strips that were transported on ice after collection. For each sample a 2-by-2 cm square of leaf tissue was used. The squares were ground with a ceramic mortar and pestle without addition of liquid nitrogen. The mashed leaf paste was then extracted using 5 mL per sample of a MeOH/$H_2O$ mixture in 2:1 ratio containing 130 ng/µL of stevioside as internal standard (but see main text for discussion of preferred internal standards). The solution was shaken thoroughly and rested at room temperature (approx. 25 °C) for 60 minutes before filtering through a filter paper (*Macherey-Nagel*, Type 615). Silicone tubing pieces of 20 mm length were prepared according to established procedures (Kallenbach *et al.*, 2014) and 60 pieces were soaked in 250 mL of extract for 1.5 hours. The silicone pieces were filtered out of the solution and dried on a fresh piece of filter paper before being stored at -20 °C, 4 °C or 30 °C.

Before the measurement, the silicone pieces were soaked in 1 mL of a MeOH:MeCN mixture in 1:1 ratio. The solution was transferred to a vial and used for the measurement with 7 µL injection volume. This method showed quite a poor extraction efficiency, exclusively extracting low polarity compounds. Due to the poor performance, it was omitted from Figure 1 and the discussion in the main manuscript.

##### 1.3. UHPLC-IM-MS/MS Measurements

The broad screening employed the following gradient at a constant flowrate of 0.6 mL/min: (i) linear increase from 5% B to 50 % B in 1.25 min; (ii) linear increase to 75% B until 1.6 min; (iii) linear increase to 100% B until 3.3 min; (iv) holding 100% B until 4.4 min (v) back to the starting conditions of 5% B until 4.5 min; (vi) equilibration for 1.1 min until the next run.

Mass spectrometry measurements were performed in "TIMS on" mode using positive ionisation. Source parameters were set as followed: End plate offset of 500 V, capillary voltage of 4500 V, nebulizer pressure of 2.2 bar ($N_2$), heated dry gas flow of 10.0 L/min with a temperature of 250 °C. Based on the recommendations for reporting IM-MS measurements (Gabelica *et al.*, 2019), the relevant TIMS parameters are as follows: a ramp time of 50 ms was chosen with an inverse reduced mobility range of 0.45–1.45 1/K0. Ion charge control (ICC) was active with a target count of 5 million. The radio frequency of the TIMS ion funnels was set at 150 Vpp. The Δ6 (ramp start → accumulation exit) voltage was set at 100.0 V. $N_2$ of at least 4.5 purity was provided by the university main supply and was used as a drift gas after purification by a HC Big Supelpure HC Hydrocarbon Trap from Sigma Aldrich (St. Louis, MO, USA). Fragmentation spectra were recorded using PASEF acquisition at 2 PASEF scans between full scans for a cycle time of 0.28 seconds.

We found that the short TIMS ramp time can cause errors including forced instrument standby, which led to an interruption of two of our measurement sequences. When adapting the method listed above, we recommend increasing the ramp time to at least 80 ms. To maintain a comparable cycle time, the number of PASEF scans can be reduced to 1 PASEF scan between full scans.

#### 2. LLE optimisation Methods

##### 2.1. Leaf Tissue Collection 2022

Maize plants of variety LG 31272 were grown under field conditions at the Strickhof competence centre in agricultural sciences (Eschikon, Switzerland, 47.4411590, 8.6811053) and tissue collection took place 14 weeks after seeding on 28$^{th}$ July 2022. The day before collection, twelve plants were tagged and five of them were sprayed with a solution containing 250 µM methyl jasmonate (MeJa) and 0.1% Tween-20 in water, while the remaining seven were sprayed with a control solution of 0.1% Tween-20 in water. From each plant, the leaves between approx. 1 and 1.8 meters above ground were sprayed with the hormone or control solution until the upper leaf side was fully covered. Four extraction solutions were prepared, all of which contained a MeOH/$H_2O$ 2:1 mixture. Two extraction solutions were spiked with stevioside (100 ng / mL) as internal standard (but see main text for discussion of preferred internal standards) and one of the spiked solutions was further acidified with 0.1% formic acid. The other two extraction solutions were spiked with camphorsulphonic acid and glycyrrhizic acid as internal standards, respectively (20 ng / mL), and no formic acid was added. Eppendorf tubes were filled with 200 µL of the extraction solutions before sample collection the following day.

From each plant, three central leaves were selected for collection. The tip (roughly 10 cm) was cut off before collecting eight leaf strips of 2 cm width for leaf tissue storage. Half of the strips were shock-frozen in liquid nitrogen and stored on dry ice, and the other half were stored in envelopes and air-dried. Following that, a 6 mm diameter hole punch (*Milian*, Vernier, Switzerland) was used to collect 12 leaf disks from the same leaf previously used for strip collection. The disks were collected into the previously prepared Eppendorf tubes, which contain 200 µL extraction solution. The tubes were thoroughly shaken before being stored in a cooling box filled with ice packs.

#### 2.2. Sample Preparation

##### 2.2.1. Frozen leaf and air-dried leaf storage samples

The shock-frozen leaf tissue was stored at -20 °C or 30 °C, while the air-dried leaf tissue was further dried to completeness at 30 °C overnight before being stored at -20 °C, 4 °C, or 30 °C.

Before the measurement, leaf pieces of 1.8 $cm^2$ were cut and placed inside microcentrifuge tubes containing metal beads. The tubes were then flash-frozen in liquid nitrogen before grinding the tissue using a TissueLyser II (*Qiagen*, Düsseldorf, Germany) until a powder was obtained. After grinding, 700 µL of the acidified extraction solution containing stevioside as internal standard was added to the tubes and samples were vortexed before adding of 500 µL of chloroform. The samples were thoroughly shaken and vortexed again and centrifuged (*Eppendorf* Centrifuge 5430 R, Hamburg, Germany) for 1 minute at 14'000 rfc. The upper phase was then transferred to vials and used for measurements with a 2 µL injection volume.

###### 2.2.2. Leaf disks in solution

The leaf disk samples were ground in solution using a micropestle (*Faust Laborbedarf*, Schaffhausen, Switzerland) attached to an electric drill (*Makita* DF333D, Kortenberg, Belgium) until a paste was obtained. After grinding, 500 µL of the appropriate extraction solution were added and the samples were thoroughly vortexed (*Thermo Scientific*, Digital Vortex Mixer, Waltham, MA, USA) followed by addition of 500 µL of chloroform. The samples were then vortexed again and then left to rest for approximately 10 minutes until phase separation was complete. 400 µL of the upper water/methanol phase were then transferred to a new Eppendorf tube and stored at -20 °C, 4 °C, or 30 °C. All extractions were completed within 30 hours of sample collection.

Before the measurement, samples were centrifuged for one minute at 12'000 rfc (*Eppendorf* 5417 R, Hamburg, Germany) and the supernatant was transferred to vials to remove remaining particles. This solution was used for measurements with a 2 µL injection volume.

##### 2.3. UHPLC-MS/MS Measurements

The LLE optimisation experiment employed the following gradient at a constant flowrate of 0.6 mL/min: (i) 5% B isocratic from 0.0 to 0.4 min; (ii) linear increase to 35% B until 2.8 min; (iii) linear increase to 75% until 3.2 min; (iv) linear increase to 100% B until 3.3 min, (v) holding 100% B until 4.4 min (vi) back to the starting conditions of 5% B until 4.5 min; (vii) equilibration for 1.1 min until the next run.

Mass spectrometry measurements were done in "TIMS off" mode and using positive and negative ionisation. The switch from "TIMS on" to "TIMS off" was done to produce higher quality fragment spectra as PASEF frequently generated spectra with fewer than 5 fragment masses being recorded. The loss of information about collisional cross section (CCS) would be significant for larger molecules (e.g., lipids or peptides), but as they are not the focal point of our study losing some information on CCS values is worthwhile when obtaining more insightful fragment spectra in return. Source parameters for positive mode were set as follows: end plate offset of 500 V, capillary voltage of 4500 V, nebulizer pressure of 2.2 bar (N2), heated dry gas flow of 10.0 L/min with a temperature of 250 °C. In negative mode the nebulizer pressure, gas flow, temperature and end plate offset are identical to positive, with the capillary voltage set to 3600 V. Fragment spectra were acquired using AutoMS/MS mode with fragmentation energies of 20 and 50 eV at a 0.3 second cycle time.

##### 2.4. Selection of internal standard compounds

As written in section 2.1, three different compounds were evaluated as internal standards. In our laboratory, camphorsulphonic acid (CSA) has been used for exploratory studies for many years (Bucher, 2017), but the rather low mass of 232 and poor ionisation in positive mode are notable downsides. We attempted to find a compound that could be used instead with the ideal properties being as follows:

- Good ionisation behaviour in positive and negative mode
- Stability over a long storage duration
- Within the mass range of 300-800 to better represent larger secondary metabolites
- Not naturally occurring in the plants we research (mainly maize)

The two compounds stevioside and glycyrrhizic acid are commercially available natural products that seemed like promising candidates in terms of ionisation behaviour and stability. This is best seen in stevioside which is sold as sweetener without cooling and with long expiry durations. Neither of the compounds is naturally present in maize and both have a mass slightly above 800, all of which then lead us to test them for this study in more detail. The results however indicate that they still show some significant downsides compared to CSA with glycyrrhizic acid showing a decreasing ion signal over the storage duration and stevioside showing the in-source fragmentation issues outlined in the main manuscript.

### 3. MetaboScape workflow

#### 3.1. Peak Extraction for broad screening (4D workflow)

Peak extraction was done using MetaboScape 2022b using the 4D workflow. All replicates were grouped together and used for the peak filter settings as follows. To be included in the final table, a signal was required to be present in at least 3 samples across the full dataset and additionally, in 75% of samples in at least one group. As there were 4 replicates, any signal that was only present in one sample group, but in at least 3 out of 4 of the replicates, remained in the final table. If a peak was present in 10 samples, but the samples were spread across 10 different sample groups, it would be filtered and not be present in the final peak table.

For peak picking, an intensity threshold of 200 was used with 80 points across the peak. For recursive peak picking, the number of points per peak was reduced to 15. Peaks were picked across the full runtime and mass range and MS/MS spectra were imported by averaging. The internal mobility and mass calibration functions were set to use the time from 0.05 min to 0.3 min as a calibration segment. Mobility calibration was performed on the signals of the tune mix (Compendium CCS variant) and mass calibration was done on sodium formate cluster signals.

#### 3.2. Peak Extraction for LLE optimisation (3D workflow)

Peak extraction was done using MetaboScape 2022b using the 3D workflow with the same filter settings as for the 4D workflow. For the peak picking an intensity threshold of 1500 was used with 7 points across the peak. For recursive peak picking, the number of points per peak was reduced to 5. Peaks were picked across the full runtime and mass range and MS/MS spectra were imported by averaging. The internal mass calibration function was set to use sodium formate cluster signals from 0.05 min to 0.35 min.

#### 3.3. Merging, blank subtraction and normalisation

The peak tables of the positive and negative polarity were merged using a mass tolerance of 3 ppm and a retention time shift tolerance of 7 seconds as a criterion to merge buckets.

Following that, signals that were detected in blank samples were removed from further analysis unless a non-blank sample had a signal intensity more than 3 times greater than the maximum blank sample. The dataset was then normalised on the intensity of the internal standard to compensate for sample handling variation. For stevioside, the selected signal was $[M+HCOO]^-$, for CSA and glycyrrhizic acid, the selected signal was $[M-H]^-$. As the broad screening was only measured in positive mode, no merging took place and for normalisation the $[M+NH_4]^+$ signal of stevioside was used.

##### 3.4. Annotation, PCA evaluation and export

All peaks were annotated using the implemented SmartFormula and MS/MS Library matching functions. For SmartFormula annotation elements CHNOPS are available with enabled element ratio filters using the “common” pre-set. The *m/z* windows are set to 1 and 3 ppm for narrow and wide annotation and the mSigma windows are set to 20 and 50 respectively. Spectral library matching employs the following narrow window parameters (wide window in brackets): 1.0 (3.0) *m/z*, 25 (1000) mSigma, 800 (250) MS/MS Score, 1 (3) % CCS. A combination of open source and commercial spectral libraries are used for the annotation and each annotation lists the library that was matched. The principal component analysis was performed inside MetaboScape using Pareto scaling. In SI Section 6, **Fig. S15** and **S16** show the normalisation effect of Pareto scaling done in MetaboAnalyst 5.0 (Pang *et al.*, 2021). The PCA data were then exported to .csv by copy-paste into Excel. Sample groups had to be re-added after the export in Excel. The export does not include the percentage of explained variance for the various principal components, so those values were then manually copied when required for figure plotting. Library annotations were exported as InChi codes in .tsv format and then used for classification using ClassyFire (Djoumbou Feunang *et al.*, 2016).

#### 4. Python workflow for figure plotting

##### 4.1. Data handling

Plotting of figures was done using the python script “PCA_Visualisation.py” (SI section 4.2) inside the Spyder IDE. The variables PC1Var and PC2Var had to be manually set by copying them from MetaboScape as they are not being exported into the .csv file using the MetaboScape export. The figures were then saved as .png files.

##### 4.2. Python script used for plotting of PCA data.

```
# -*- coding: utf-8 -*-
"""
Created on Fri Oct 22 07:42:26 2021

@author: jlang
"""
#%% Import
import pandas as pd
import numpy as np
#import math
from bokeh.plotting import figure, output_file, show
from bokeh.models import ColumnDataSource, Ellipse
from bokeh.transform import factor_cmap, factor_mark
import os
from scipy.stats import chi2

#%% calculation of a confidence ellipse, coded in part by ChatGPT
def confidence_ellipse(x, y, color, p):
    # Calculate the mean and covariance of the data
    mean = np.array([np.mean(x), np.mean(y)])
    cov = np.cov(x, y)

    # Calculate the eigenvalues and eigenvectors of the covariance matrix
    eigenvalues, eigenvectors = np.linalg.eig(cov)

    # Sort the eigenvalues and eigenvectors in descending order
    sorted_indices = np.argsort(eigenvalues)[::-1]
    eigenvalues = eigenvalues[sorted_indices]
    eigenvectors = eigenvectors[:, sorted_indices]

    # Calculate the angle of the ellipse
    angle = np.degrees(np.arctan2(*eigenvectors[:, 0][::-1]))

    # Calculate the width and height of the ellipse
    width = 2 * np.sqrt(chi2.ppf(0.9747, 2)) * np.sqrt(eigenvalues[0])
    height = 2 * np.sqrt(chi2.ppf(0.9747, 2)) * np.sqrt(eigenvalues[1])

    # Add the confidence ellipse to the plot
    p.ellipse(x=mean[0], y=mean[1], width=width, height=height, angle=angle,
            angle_units='deg', fill_alpha=0, line_color=color, line_width=3)

    return None

#%% PC variations are hardcoded and need to be manually adapted. Sorry for that
PC1Var = 20.0
PC2Var = 16.3

#%% Import of csv file and data cleaning. csv_files allows quick selection of input data
currdir = os.getcwd()
```

```python
    all_files = os.listdir(currdir)
    csv_files = list(filter(lambda f: f.endswith('.csv'), all_files))
    csvPath = currdir + "\\" + csv_files[0]

    df = pd.read_csv(csvPath, sep=";")
    df["Time"]=df["Time"].values.astype('str')
    mask = (df['Time']=='1')
    notMask = (df['Time']!='1')
    df.loc[mask, "Time"] = df["Time"].astype('str')+" Day"
    df.loc[notMask, "Time"] = df["Time"].astype('str')+" Days"


    #%% categories in my dataset and colourmaps/symbols used in the plot
    markers = ['hex', 'square', 'triangle','cross']
    method = ['LLE', 'SPE', 'ST']
    methods = ['Extract', 'AirDry', 'LN2']
    storage = ['1 Day', '6 Days', '14 Days', '28 Days', '42 Days', '56 Days']

    # color palette by Wong (Nature Methods, 2011) tuned for color blindness
    col = ['#E69F00','#56B4E9','#009E73','#F0E442','#0072B2', '#CC79A7']
    temp = ['-20 °C', '4 °C', '30 °C', '-80 °C']
    acid = ['FormicAcid', 'NoAcid']
    stress = ['Stessed', 'Not']


    #%% general plot setup with title and labels
    output_file("PCA_scatter.html")
    plot = figure(width=800, height=650,
                  title = "Liquid-Liquid Extract Comparison - Without RT")
    plot.title.text_font_size = '18pt'
    plot.xaxis.axis_label = 'PC 1 ({} %)'.format(PC1Var)
    plot.yaxis.axis_label = 'PC 2 ({} %)'.format(PC2Var)
    plot.xaxis.axis_label_text_font_size = '12pt'
    plot.yaxis.axis_label_text_font_size = '12pt'

    #%% plot the datapoints and the confidence ellipse by category
    # Note that the iteration needs to be done by groups used for the ellipse, here methods was used
    for k in methods:
        filDF = df[df['Method'] == k]
        source = ColumnDataSource(filDF)
        plot.scatter(x='PC 1', y='PC 2', source=source, legend_label=k,
                     size = 16, fill_alpha = 0.4,
                     marker=factor_mark("Method", markers, methods),
                     color=factor_cmap("Method", col, methods))
        confidence_ellipse(filDF["PC 1"], filDF["PC 2"], "#D55E00", plot)

    plot.legend.label_text_font_size = '14pt'
    plot.legend.glyph_height = 30
    plot.legend.glyph_width = 30
    plot.legend
    plot.legend.location = "top_right"

    show(plot)
```

#### 5. R Script of MANOVA and pairwise comparison with output

```
# DATA, BASED ON SCORES

pca  <-read.csv("/Users/sramos/Desktop/data/JK.csv")

pca$method <- as.factor(pca$method)

#  LEAVING ONLY THE NUMERIC VARIABLES
pc1 <- pca[,c(-1,-2,-3,-4,-5,-6)]

# PCA ON THE pc1 MATRIX, ONLY NUMERIC VARIABLES
pca_res <- prcomp(pc1, scale. = TRUE)

# PLOTTING WITH PAM # https://www.rdocumentation.org/packages/cluster/versions/2.1.4/topics/pam

library(cluster)
autoplot(pam(pc1[-5], 3), frame = TRUE, frame.type = 'norm')

#   MANOVA ON THE PC'S AS A LINEAR COMBINATION OF VARIABLES IN FUNTION OF THE FACTOR "METHOD"
#   THE NUMBER OF PC (1-10) INDICATED UP TO WHICH PC THERE WERE SIGNIFICANT PCS, IN THIS CASE, UP TO PC5

res.man <- manova(cbind(PC1, PC2, PC3, PC4, PC5, PC6, PC7, PC8, PC9, PC10) ~ method, data = pca)
summary(res.man)      # METHOD HAS A SIGNIFICANT EFFECT, NO SURPRISE

#            Df  Pillai approx  F num  Df den  Df    Pr(>F)
#  method     2   1.8991       691     20      734   < 2.2e-16 ***
#  Residuals  375


summary.aov(res.man)  # HERE WE SEE THE THAT ONLY UP TO PC5 IS SIGNIFICANT
#Response PC1 :
#           Df    Sum Sq   Mean Sq F value    Pr(>F)
#method       2 1602700098 801350049  1667.1 < 2.2e-16 ***
#  Residuals   375  180257663    480687
#
#Response PC2 :
#           Df    Sum Sq   Mean Sq F value    Pr(>F)
#method       2 1077335749 538667874  532.72 < 2.2e-16 ***
# Residuals   375  379184656   1011159
#Response PC3 :
#           Df   Sum Sq  Mean Sq F value    Pr(>F)
#method       2 162144387 81072193  39.473 2.771e-16 ***
#  Residuals   375 770205296  2053881
#Response PC4 :
#           Df   Sum Sq Mean Sq F value  Pr(>F)
#method       2  10102678 5051339  4.0982 0.01735 *
#Response PC5 :
#           Df   Sum Sq Mean Sq F value    Pr(>F)
#method       2  13352214 6676107  7.2809 0.0007903 ***
```

```
#  Residuals   375 343850789  916935
#Response PC6 :
#           Df    Sum Sq Mean Sq F value Pr(>F)
#method       2    320300  160150  0.2148 0.8068
#Residuals   375 279563492  745503
#Response PC7 :
#           Df    Sum Sq Mean Sq F value Pr(>F)
#method       2   3711871 1855936  2.7122 0.0677 .
#Residuals   375 256612814  684301
#Response PC8 :
#           Df    Sum Sq Mean Sq F value Pr(>F)
#method       2    506816  253408  0.4834 0.6171
#Residuals   375 196582786  524221
#Response PC9 :
#           Df    Sum Sq Mean Sq F value Pr(>F)
#method       2   1719735  859868  1.7423 0.1765
#Residuals   375 185067040  493512
#Response PC10 :
#           Df    Sum Sq Mean Sq F value Pr(>F)
#method       2    109807   54904  0.1334 0.8751
#Residuals   375 154326389  411537




# RUNNING THE POSTHOC CONTRAST OF FACTOR "METHOD" TO SEE WHICH METHODS DIFFER FROM EACH OTHER. THE RESPONSE VARIABLE IS AGAIN
# A LINEAR COMBINATION OF VARIABLES OF PC 1-5.
emmeans((lm(cbind(PC1, PC2, PC3, PC4, PC5) ~ method, data=pca)), pairwise ~ method, contrast="pairwise", p.adjust="bonferroni") # it worked

#$emmeans
#method  emmean   SE  df lower.CL upper.CL
#AirDry   689.0 61.0 375      569    809.0
#Extract -325.6 43.1 375     -410   -240.8
#LN2      -40.2 63.0 375     -164     83.6
#
#Results are averaged over the levels of: rep.meas
#Confidence level used: 0.95
#
#$contrasts                                THIS IS THE IMPORTANT BIT OF THE RESULTS, THE PAIRWISE CONTRASTS
#contrast         estimate   SE  df t.ratio p.value
#AirDry - Extract     1015 74.7 375  13.580  <.0001
#AirDry - LN2          729 87.7 375   8.315  <.0001
#Extract - LN2        -285 76.4 375  -3.738  0.0006

#Results are averaged over the levels of: rep.meas
#P value adjustment: tukey method for comparing a family of 3 estimates
```

### 6. Additional figures and photographs

#### 6.1. Normalisation, Stability and Homogenisation

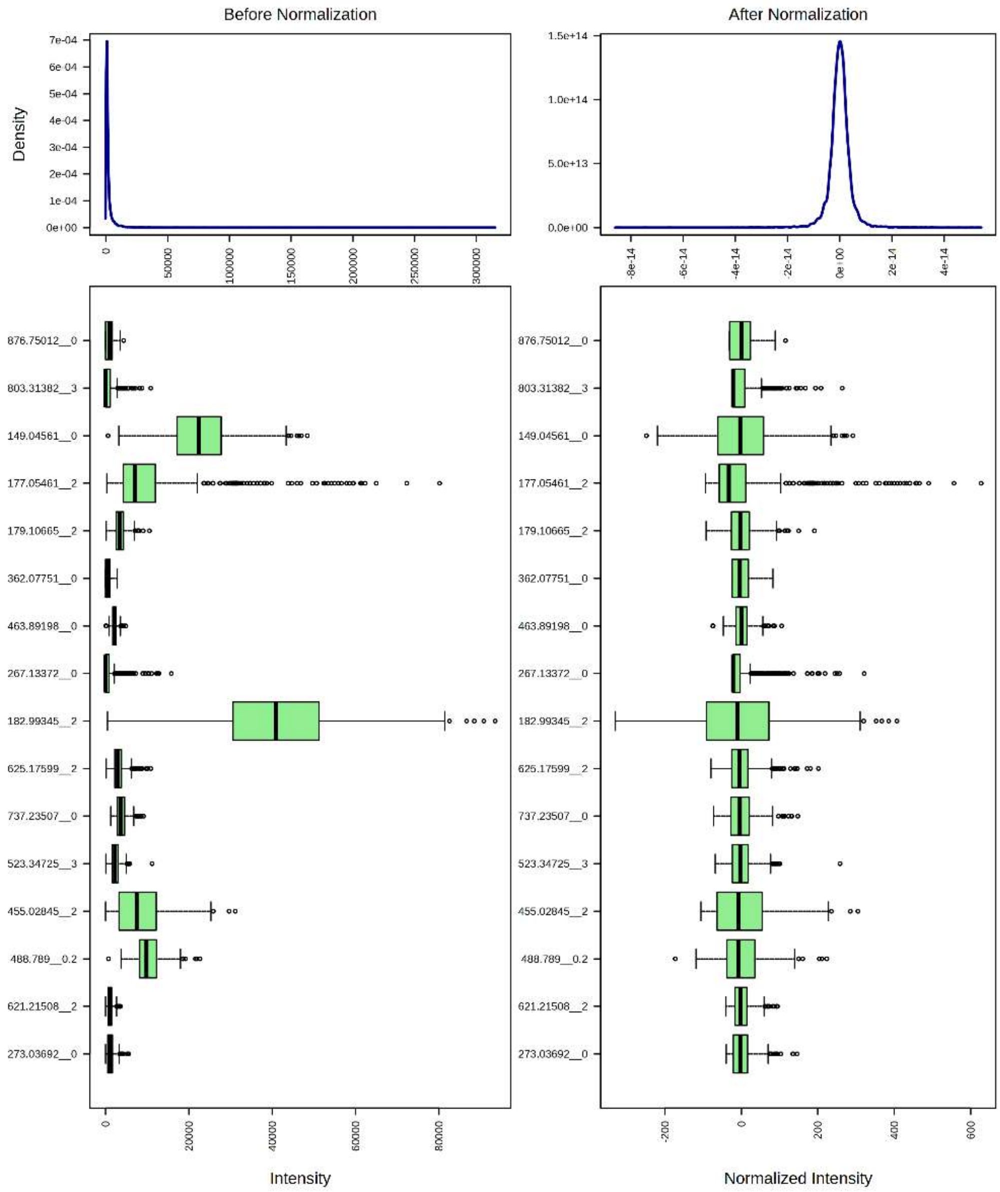


***Fig. S1:*** *Influence of pareto scaling on the peaks. Overall distribution shown on top with examples of individual peaks below.*

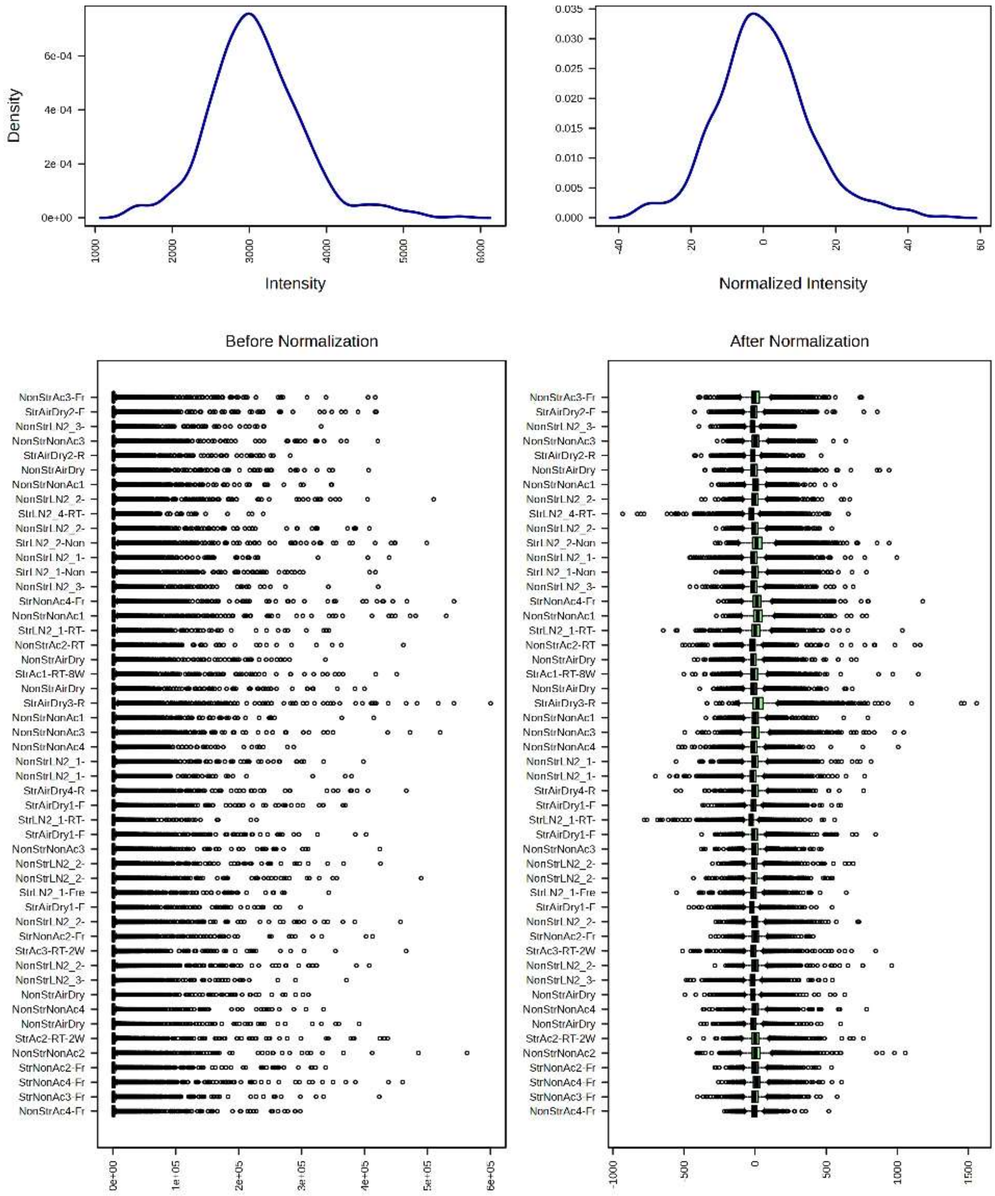


***Fig. S2:** Influence of pareto scaling on the samples. Overall distribution on the top with individual examples below.*

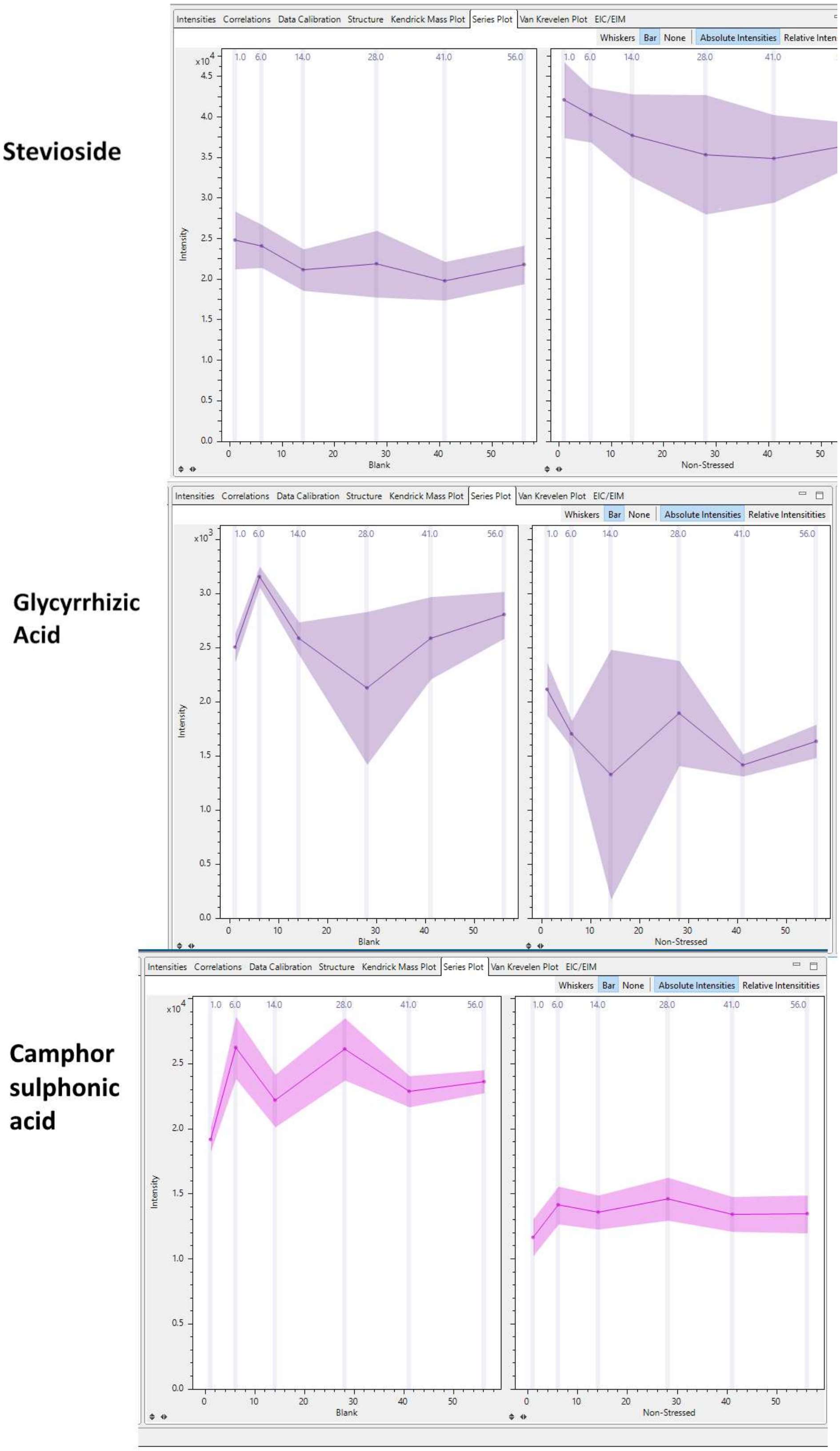


***Fig. S3:*** *Comparison of the internal standards which were assessed for stability. Shown on the left side are blank measurement of each extraction solution, the right side shows samples where a full extraction procedure was performed with the three solutions. Both stevioside and glycyrrhizic acid show a slight downwards trend while CSA is stable across the 6 time points.*

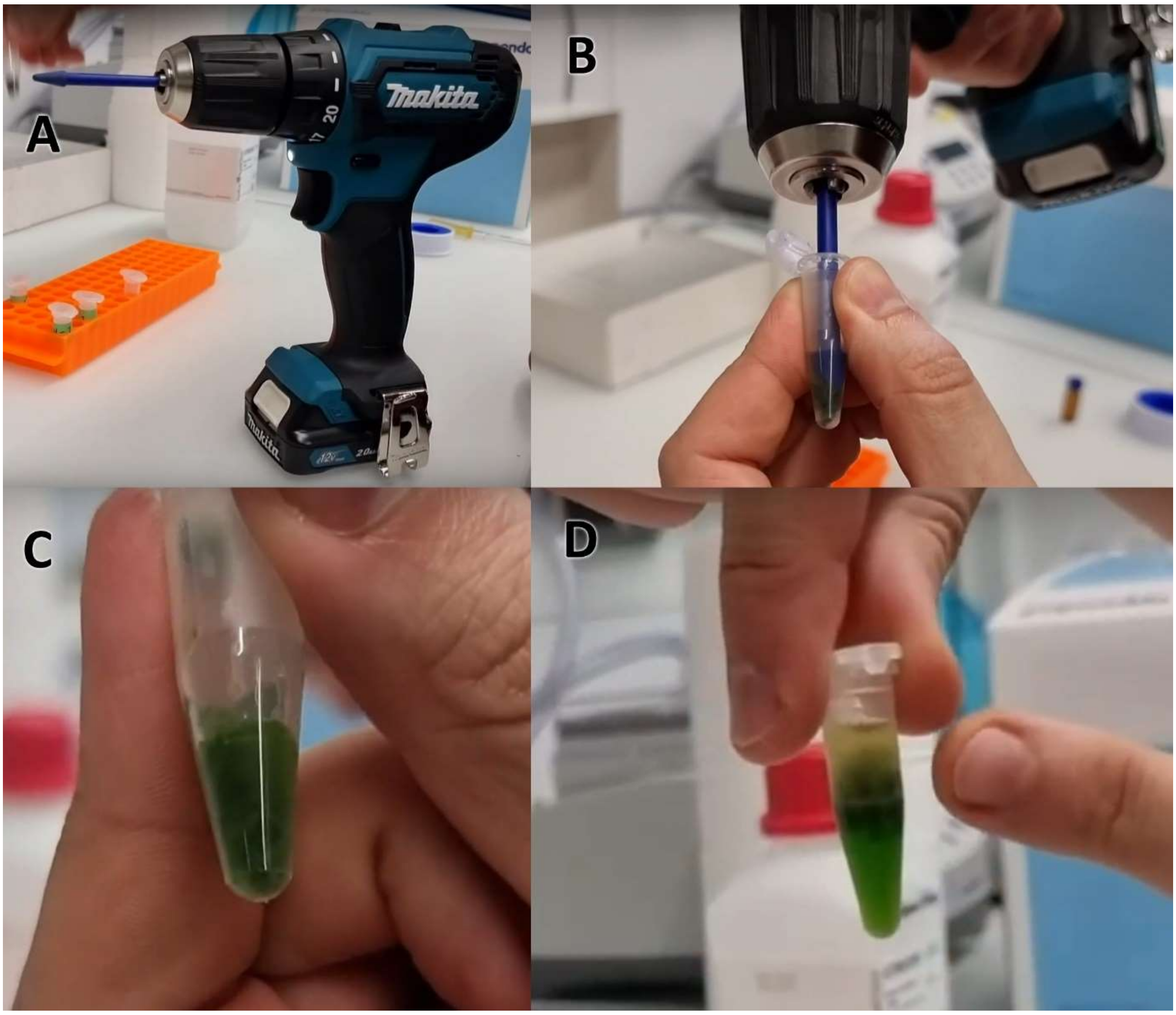


***Fig. S4:*** *Photos of the proposed on-site extraction method with a focus on homogenisation. The mobile homogenisation setup (A) is used to grind leaf tissue inside 1.5 mL microcentrifuge tubes (B) until a paste-like consistency is reached (C), which can then be used for a liquid-liquid extraction (D). All images are screenshots from our video tutorial: https://youtu.be/DpEHiu9cFdU*

#### 6.2. PCA Plots of broad screening

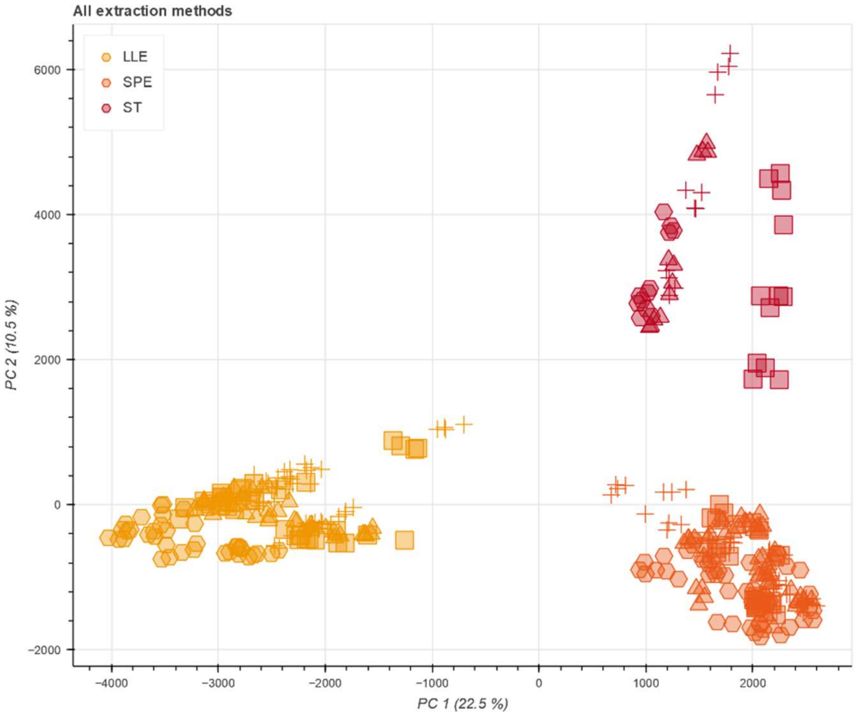


***Fig. S5:*** *PCA plot of all samples in the broad screening. The LLE, SPE and silicone tubing approaches cluster together, and evaluation of storage temperature and duration is impractical. Icons indicate storage duration; Hexagon = 1 Day, Triangle = 1 Week, Square = 1 Month, Cross = 75 Days.*

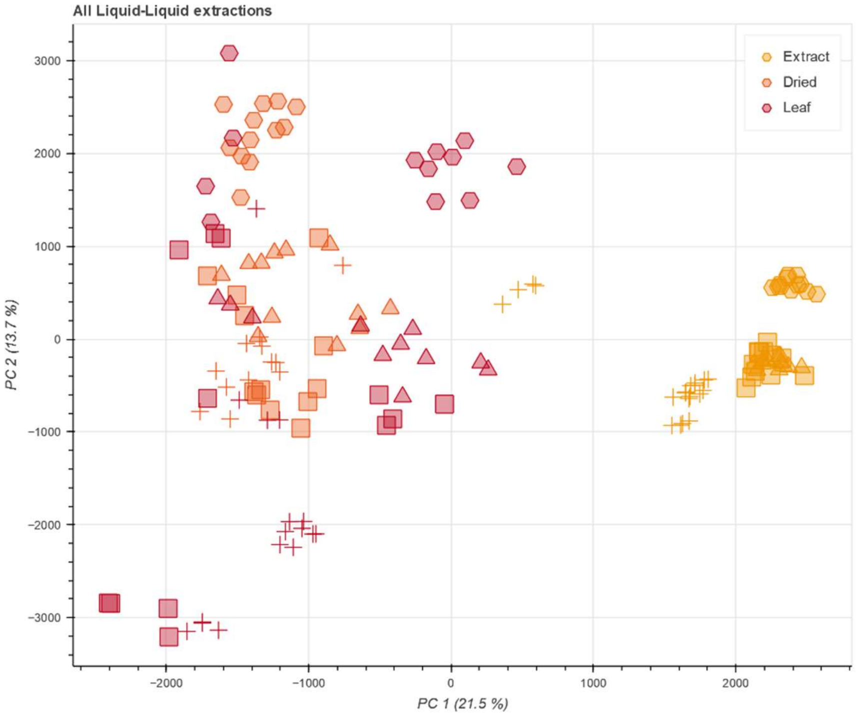


***Fig. S6:*** *PCA plot of all samples prepared by LLE. Note that the grinding of leaf tissue for the extract storage was unoptimized leading to the large differentiation of Extract storage and the leaf storage. Secondly, the extract storage showed a much better reproducibility (tighter grouping) and reduced impact of storage duration. Icons indicate storage duration; Hexagon = 1 Day, Triangle = 1 Week, Square = 1 Month, Cross = 75 Days.*

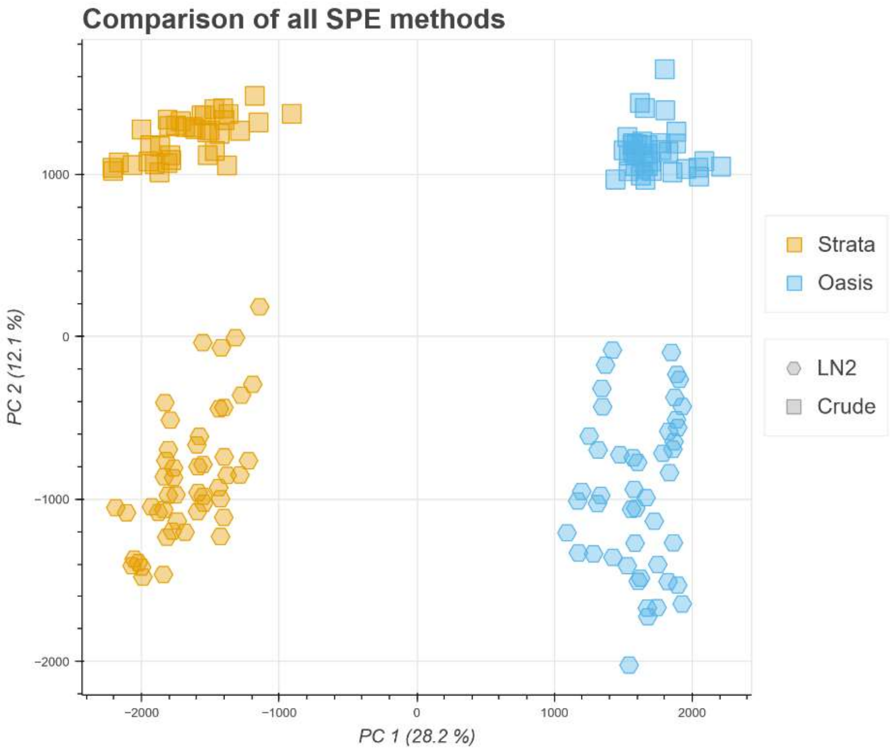


***Fig. S7:*** *PCA plot of all samples prepared by SPE with a clear separation based on the SPE brand (color) and a secondary separation based on the homogenisation (hexagon = pulverisation with liquid nitrogen, square = homogenisation at ambient temperature).*

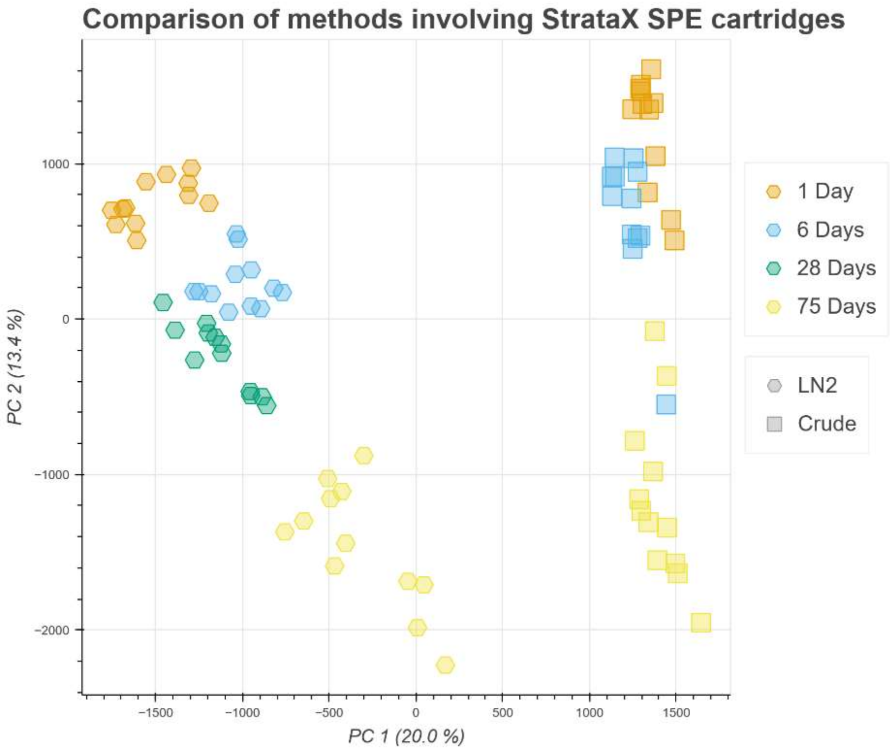


***Fig. S8:*** *PCA plot of all methods done with StrataX Pro SPE cartridges. The left side contains all samples stored as extract which underwent SPE directly before measurement (FE-Fil), the right side contains the samples stored on the SPE cartridge (CE-Stra).*

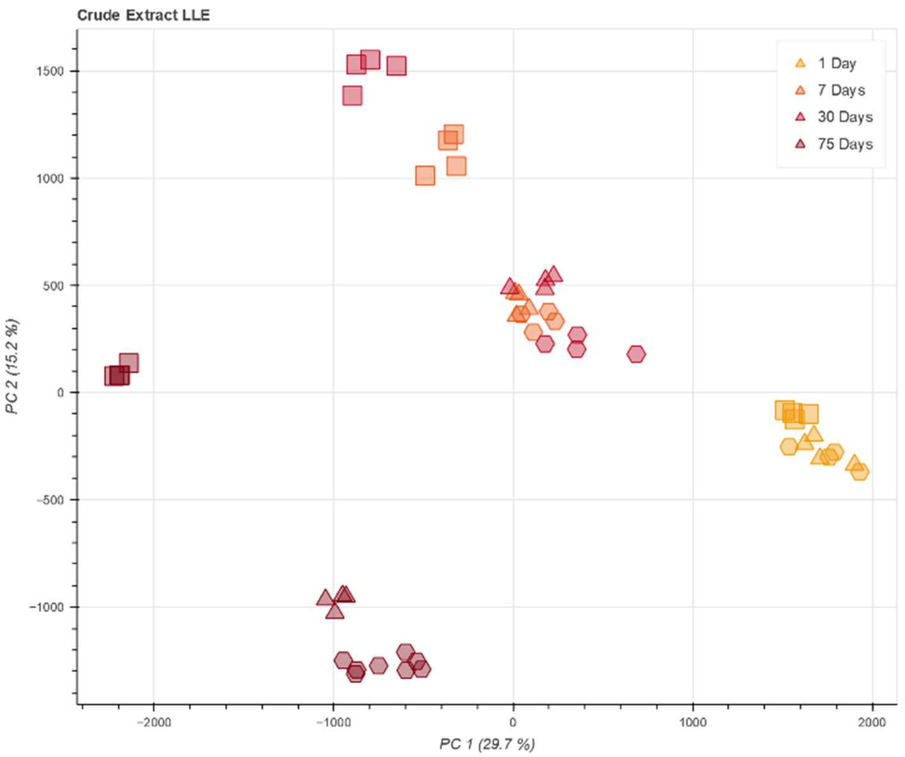


***Fig. S9:*** *PCA plot of the extract storage after LLE (CE-LLE) over time. Colour indicates storage duration, icons indicate storage temperature (hex = -20 °C, triange = 4 °C, square = 30 °C).*

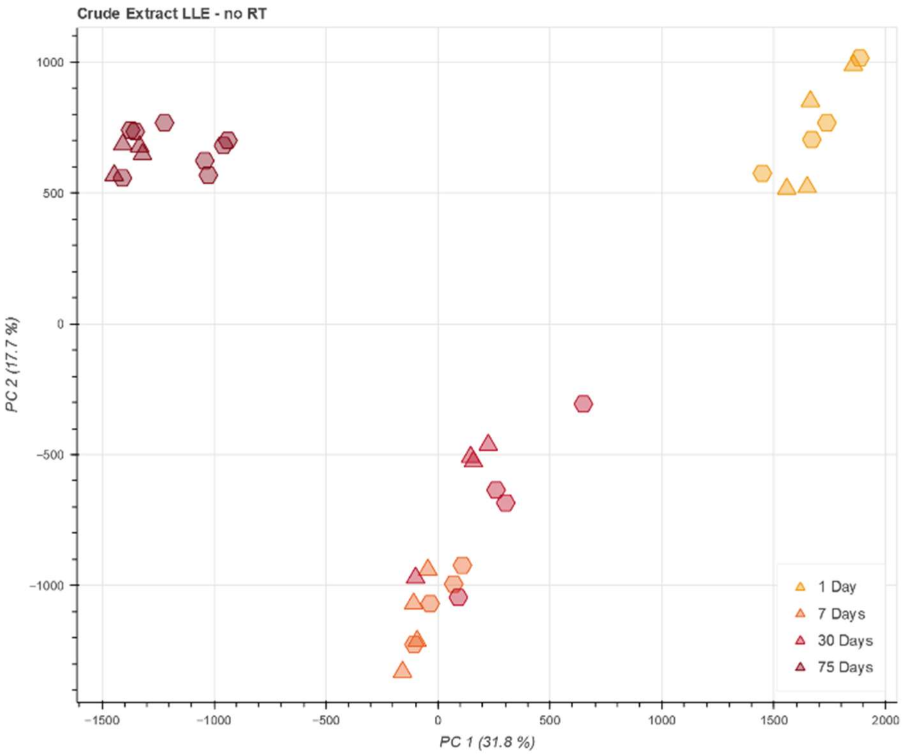


***Fig. S10:*** *PCA plot of the extract storage after LLE (CE-LLE) over time, excluding samples stored at room temperature (30 °C). Colour indicates storage duration, icons indicate storage temperature (hex = -20 °C, triange = 4 °C).*

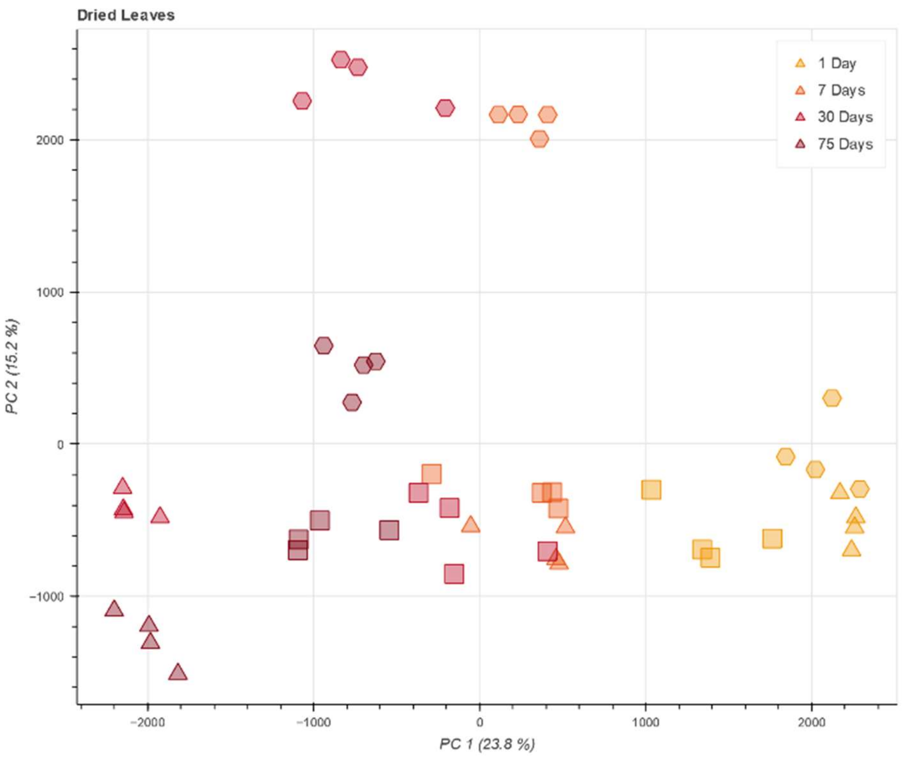


***Fig. S11:*** *PCA plot of air-dried leaf storage. Notably, the samples stored in a freezer seemed to be the outliers here (hexagons which separate out on PC 2).*

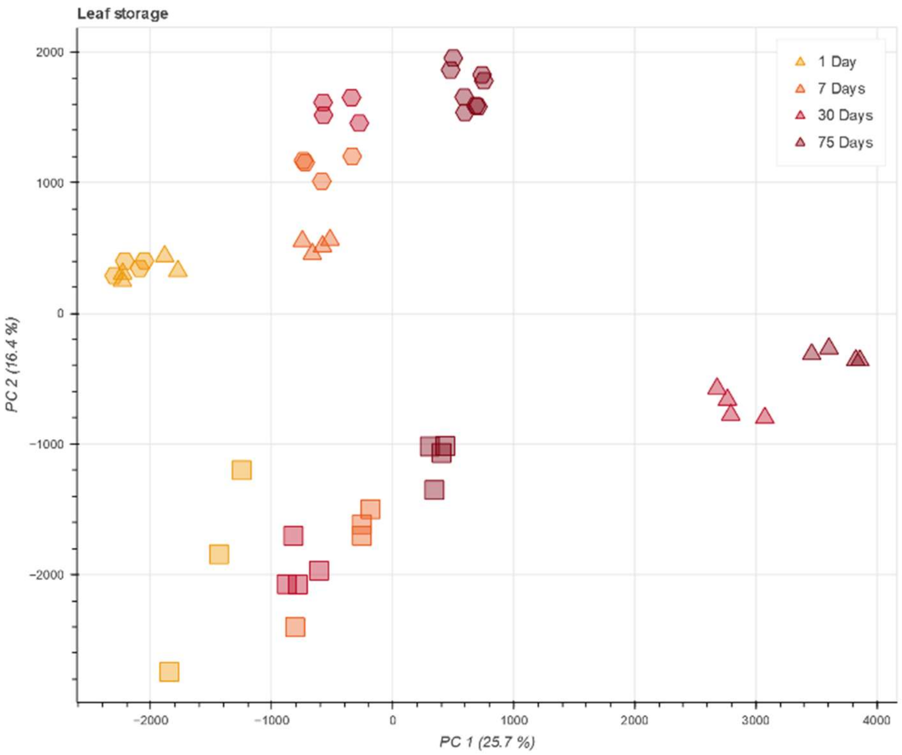


***Fig. S12:*** *PCA plot of shock-frozen leaf tissue. Note the far separation of storage at 4 °C (triangles on the right) which was caused by fungal growth on the leaf, and the separation of samples stored at 30 °C (squares) which are very similar to air-dried samples.*

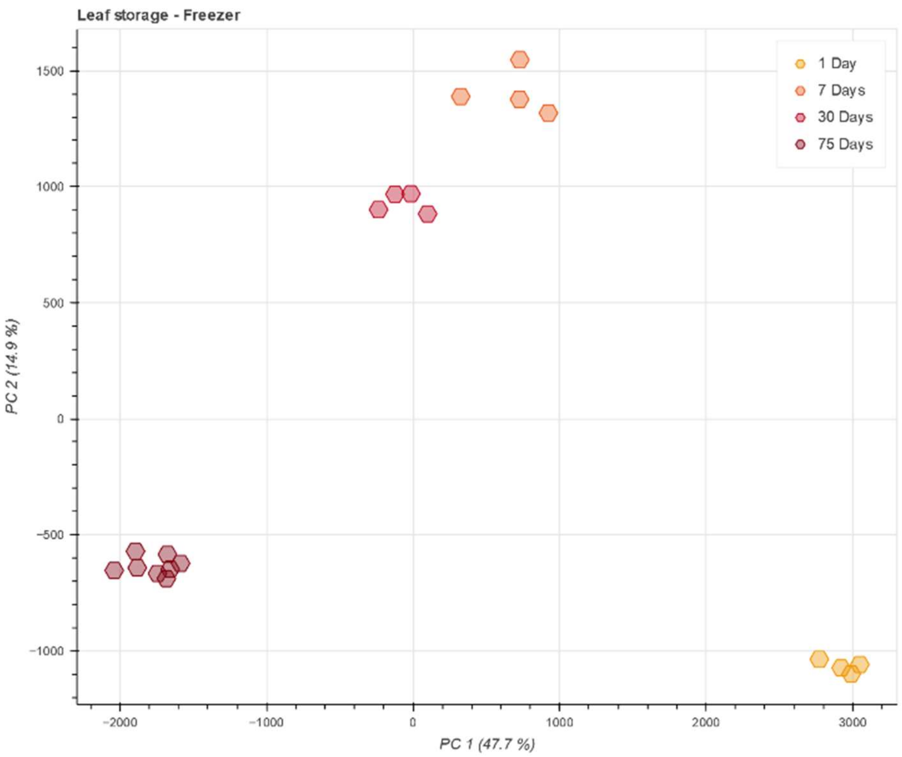


***Fig. S13:*** *PCA plot of shock-frozen leaf storage at -20 °C. Small differences are observed between 7 and 30 days, larger gaps are present between the other time points.*

#### 6.3. PCA Plots of LLE optimisation

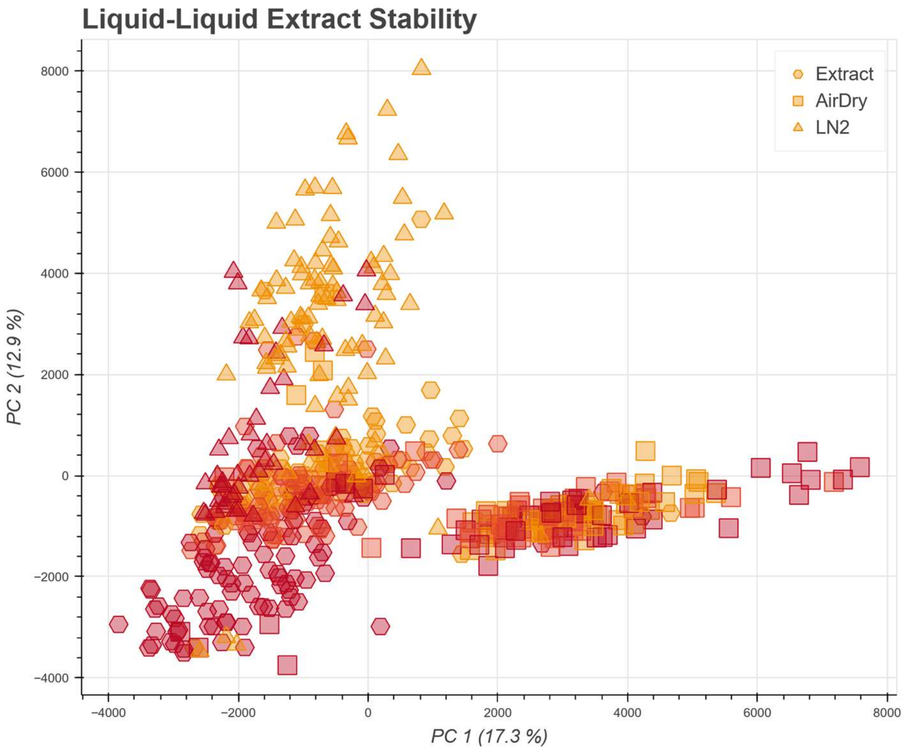


***Fig. S14:*** *PCA of all samples of the LLE optimisation. Temperature indicated by colour, the darker the warmer.*

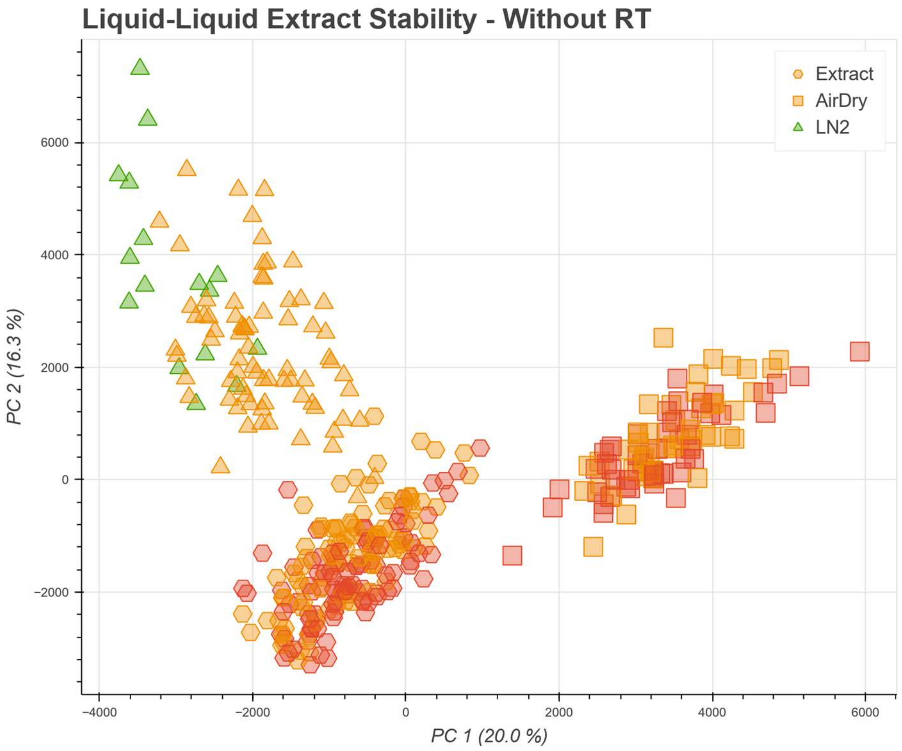


*__Fig. S15:__ PCA of LLE optimisation samples excluding samples stored at 30 °C. The colour indicates storage temperature, yellow = -20 °C, red = 4 °C, green highlights shock-frozen leaf tissue, measured after a day at -20 °C as the ideal case.*

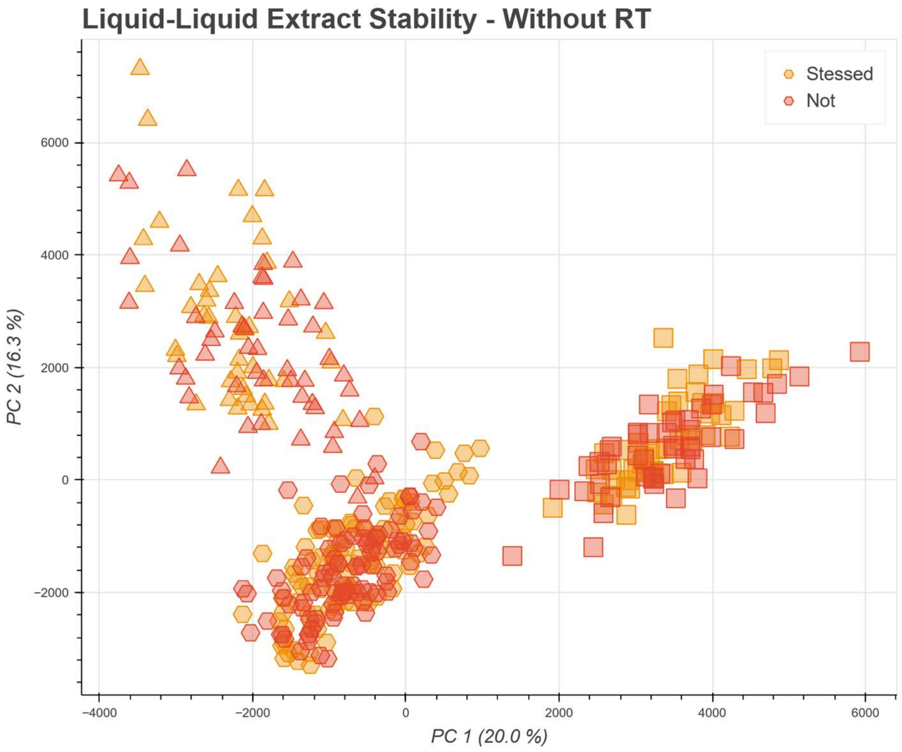


*__Fig. S16:__ Same PCA as above with different colour indication. Colour now refers to the treatment with methyl jasmonate to induce stress in the plants. No clear effect could be observed.*

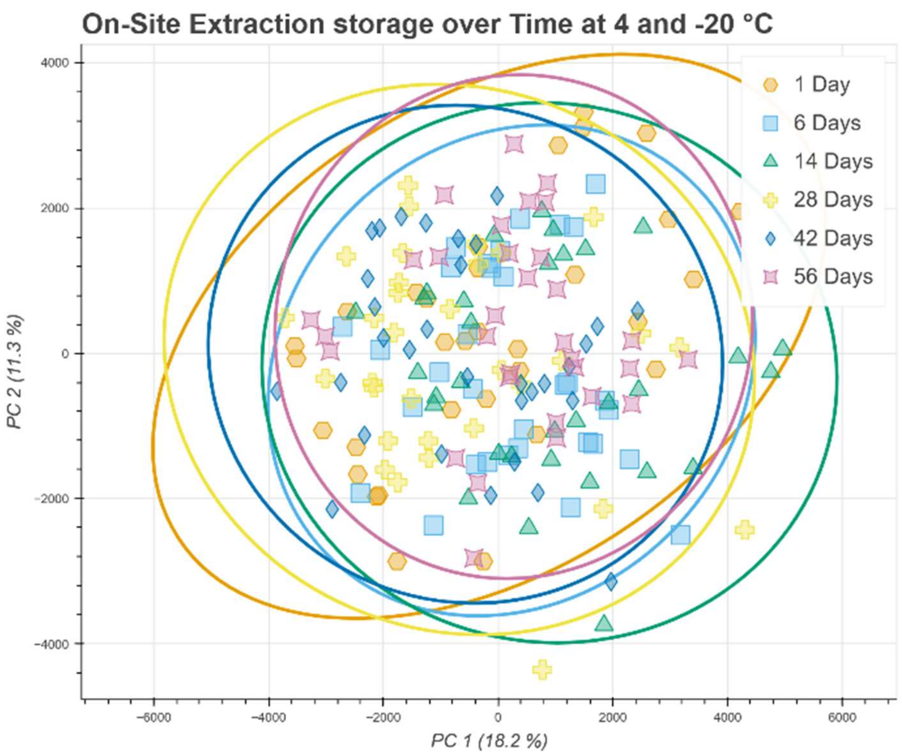


***Fig. S17:*** *PCA plot of all on-site extraction samples stored at reduced temperature. PC1 and PC2 do not contain any trends of degradation over time. The minor observed trend can only be found in PC3 and PC4 (see main text).*

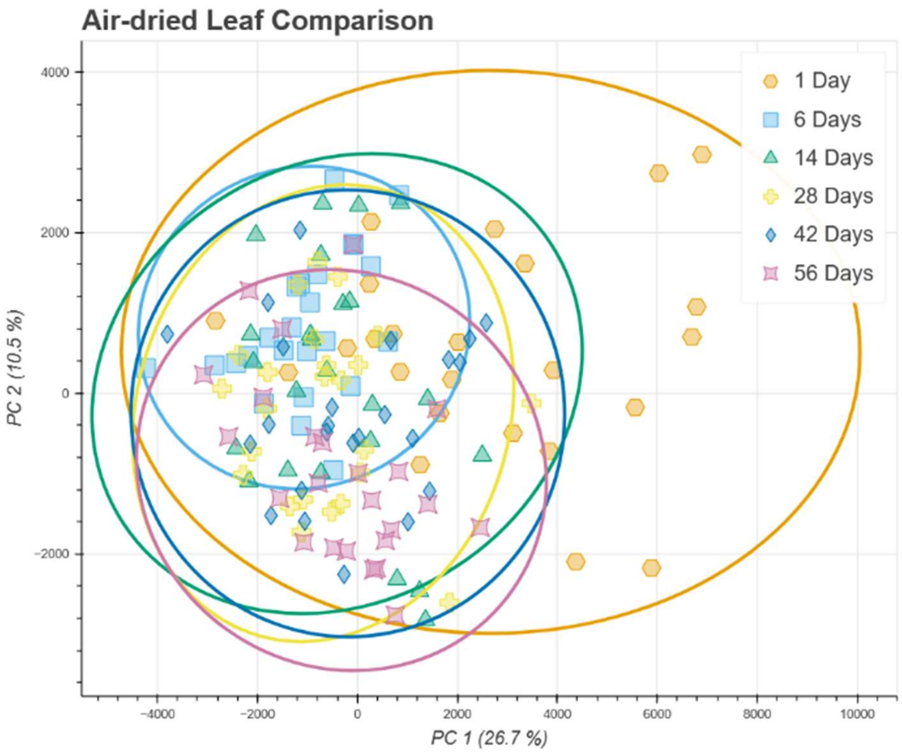


***Fig. S18:*** *PCA plot of all air-dried samples. After a minor shift from day 1 to day 6, the samples show no further shifts. It is possible that the drying process was not completed before the extraction of the samples began.*

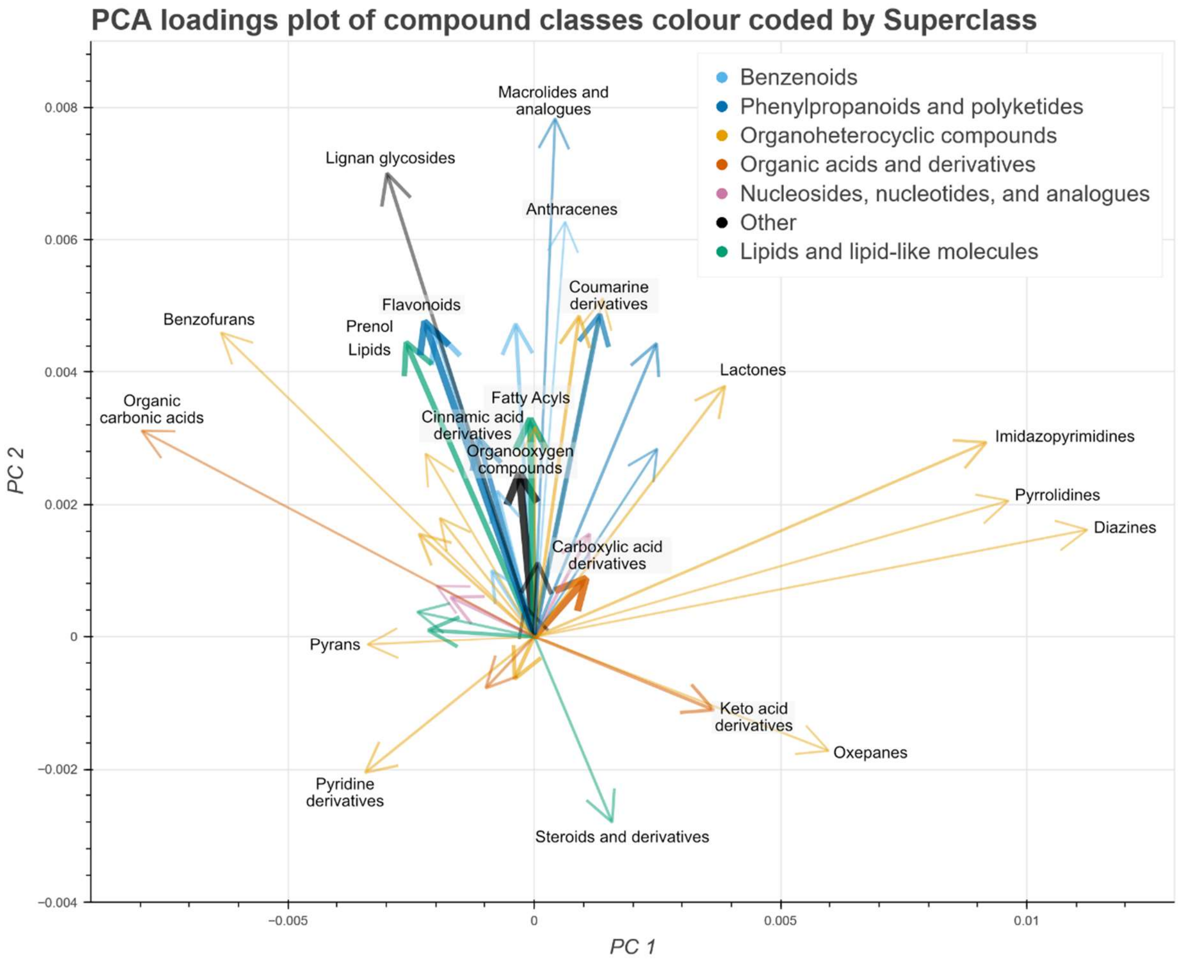


***Fig. S19:*** *Full size version of Fig. 5B for better readability of arrow labels.*

**PCA dimensions 1 to 5 of the LLE Optimisation experiment**

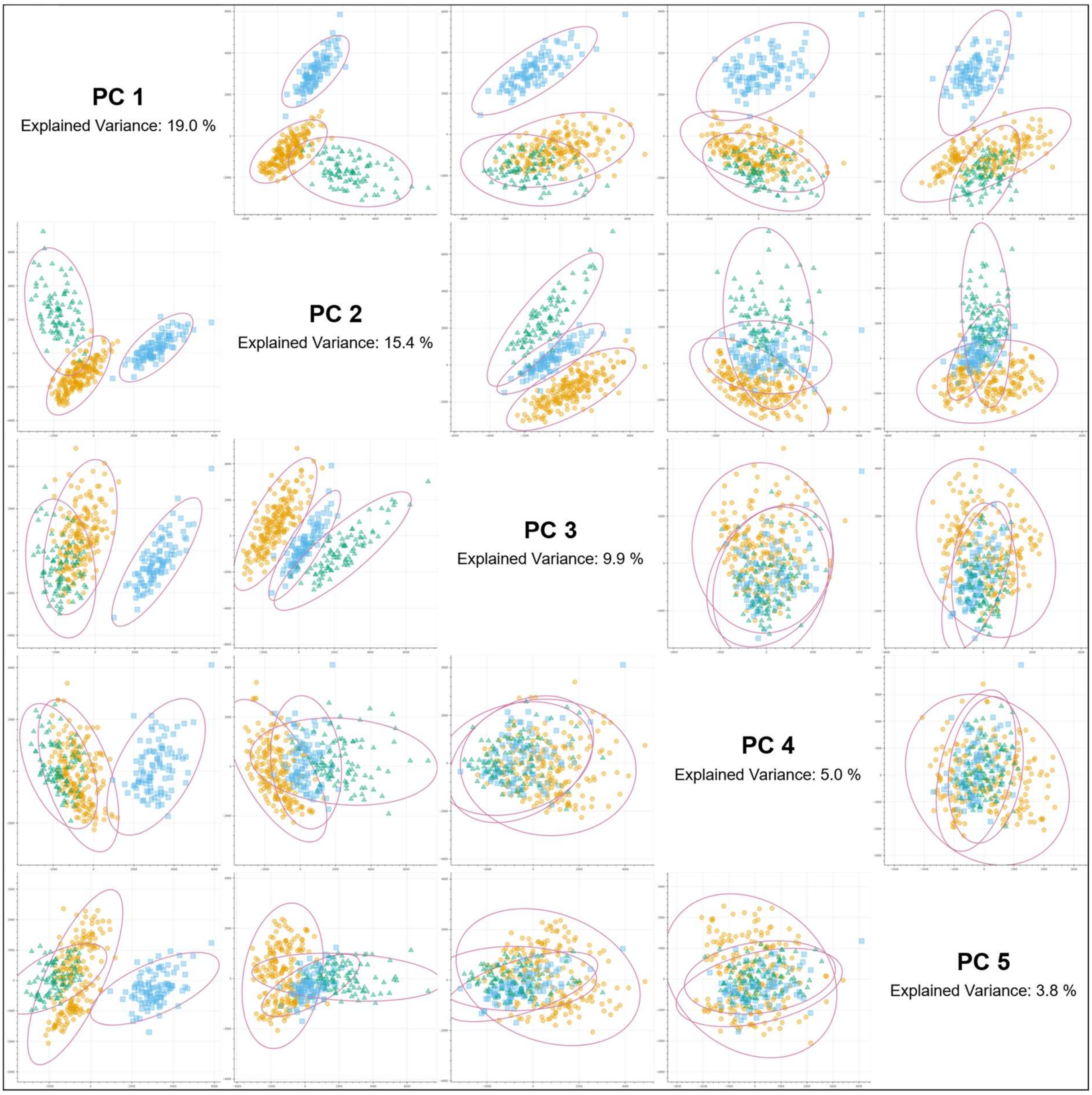


***Fig. S20:*** *Grid plot showing the PC1 to PC5 dimensions against each other for the full LLE optimisation dataset (see also Fig. 5A, which is just the PC1 against PC2 plot in this figure. According to the MANOVA analysis, dimensions 1, 2, 3 and 5 show highly significant separation of the groups with PC4 being barely significant.*

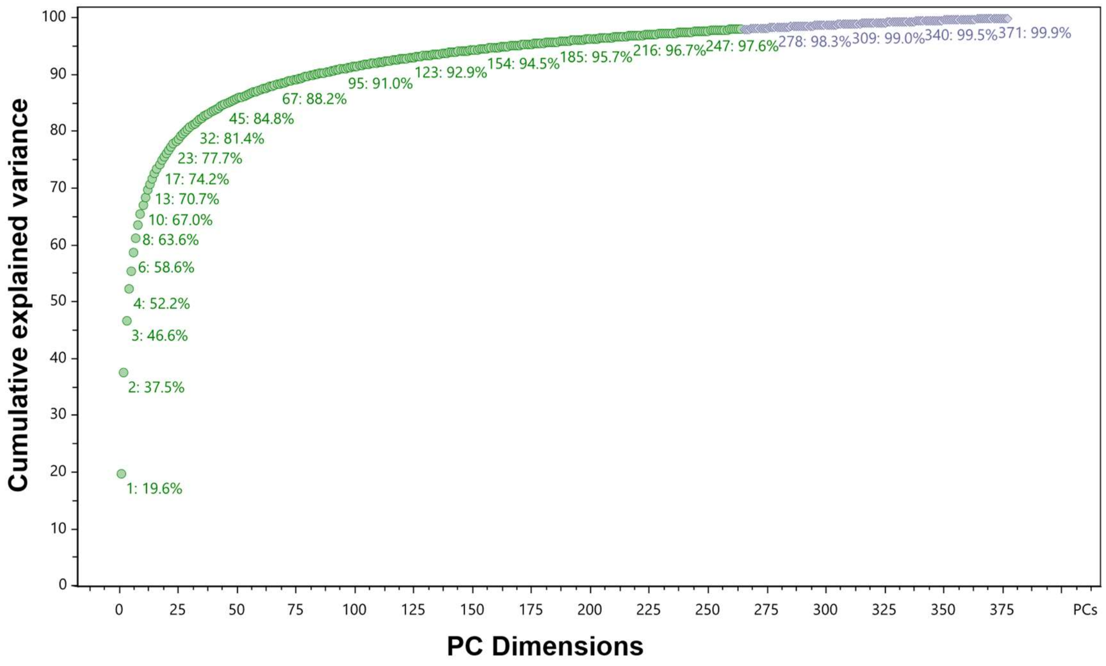


***Fig S21:*** *Cumulative variance of the PC dimensions of Fig 5A. The green colour indicates the dimensions that would be required to explain 98% of the total variance.*
